# Promoter-adjacent DNA hypermethylation can downmodulate gene expression: *TBX15* in the muscle lineage

**DOI:** 10.1101/2022.11.14.516527

**Authors:** Kenneth C. Ehrlich, Michelle Lacey, Carl Baribault, Sagnik Sen, Pierre Olivier Esteve, Sriharsa Pradhan, Melanie Ehrlich

## Abstract

*TBX15*, which encodes a differentiation-related transcription factor, displays promoter-adjacent DNA hypermethylation in myoblasts and skeletal muscle (psoas) that is absent from non-expressing cells in other lineages. By whole-genome bisulfite sequencing (WGBS) and enzymatic methyl-seq (EM-seq), these hypermethylated regions were found to border both sides of a constitutively unmethylated promoter. To understand the functionality of this DNA hypermethylation, we cloned the differentially methylated sequences (DMRs) in CpG-free reporter vectors and tested them for promoter or enhancer activity upon transient transfection. These cloned regions exhibited strong promoter activity and, when placed upstream of a weak promoter, strong enhancer activity specifically in myoblast host cells. *In vitro* CpG methylation targeted to the DMR sequences in the plasmids resulted in 86 - 100% loss of promoter or enhancer activity, depending on the insert sequence. These results as well as chromatin epigenetic and transcription profiles for this gene in various cell types support the hypothesis that DNA hypermethylation immediately upstream and downstream of the unmethylated promoter region suppresses enhancer/extended promoter activity, thereby downmodulating, but not silencing, expression in myoblasts and certain kinds of skeletal muscle. This promoter-border hypermethylation is not found in cell types with a silent *TBX15* gene probably because they have no transcription to modulate. *TBX18, TBX2, TBX3* and *TBX1* display *TBX15*-like hypermethylated DMRs at their promoter borders and preferential expression in myoblasts. Therefore, promoter-adjacent DNA hypermethylation for downmodulating transcription to prevent overexpression may be used more frequently for transcription regulation than currently appreciated.

## 1. Introduction

The transcription factors (TFs) that play key roles during embryonic differentiation often also have important tissue-specific postnatal functions [1]. *TBX15* encodes a TF in the T-box family involved in embryogenesis, regulation of various postnatal developmental processes, and homeostasis [2–6]. T-box domain transcription factors can act as repressors and, less frequently, as activators of transcription [3]. Inactivation of these TFs by inherited mutations in humans leads to congenital defects in skeletal/craniofacial bone formation (TBX15, TBX18, TBX1, TBX3, TBX4, TBX5, TBX6, TBX22, and TBXT proteins) and heart formation (TBX1, TBX3, TBX5, and TBX20 proteins) [7,8]. Studies of animal models and human cell cultures indicate additional functions of T-box TFs, such as regulation of skin pigmentation (murine Tbx15), lung or liver hypoplasia (human TBX2 and TBX3), early embryogenesis (murine Tbr2 and Tbxt), and skeletal muscle (SkM) physiology (murine Tbx15, frog/bird-specific Tbx16, and *C. elegans* TBX-2)[3,9–12]. Studies of mice indicated a strong association of *Tbx15* expression with muscle fiber type [10] and a positive association of *Tbx15* expression with lean mass and several metabolic phenotypes [13].

Evidence for the importance of *TBX15* to the human SkM lineage was seen in our previous comparison of transcriptomic and epigenomic profiles of *Tbx15/TBX15* in SkM tissue and myoblasts [14,15]. Myoblasts are SkM progenitor cells involved in embryogenesis and postnatal repair of muscle damage [16]. We found that human myoblast primary cultures and their differentiation product, multinucleated myotubes, express moderate levels of *TBX15* RNA while there is little or no expression of this gene in five diverse cell cultures that are not derived from mesoderm. Moreover, SkM had the highest expression of *TBX15* in a comparison of 52 studied human tissues [14,17].

Analysis of methylomes generated by reduced representation bisulfite sequencing (RRBS) surprisingly revealed that transcriptionally active *TBX15* is strongly hypermethylated both immediately upstream and downstream of the unmethylated promoter region in myoblasts and myotubes compared with 15 types of cell cultures not expressing the gene. Association of intragenic DNA hypermethylation with actively transcribed gene bodies is a frequent, but not universal, finding that could be due to gene-body DNA methylation modulating the rate of movement of the transcription complex, regulating alternative splicing, or repressing cryptic promoters, retrotransposons, enhancers, and silencers [18]. However, gene-body DNA methylation that is positively correlated with transcription is observed most strongly in DNA sequences considerably downstream of the transcription start site (TSS) [18,19], unlike the promoter-adjacent intragenic hypermethylation that we observed in *TBX15* in the SkM lineage [14].

To elucidate the role of the DNA hypermethylation around the constitutively unmethylated *TBX15* promoter in *TBX15*-expressing myoblasts and SkM, we cloned DNA sequences from several of these differentially methylated regions (DMRs) and tested their ability to act as promoters or enhancers in reporter gene transfection assays with or without *in vitro* CpG methylation targeted to these sequences. We found that the hypermethylated DNA sequences immediately upstream and downstream of the promoter have strong promoter or enhancer activity when unmethylated and transfected into myoblasts. These results and our in-depth analysis of epigenomic *vs*. transcriptomic profiles of many cell and tissue types support the hypothesis that the myoblast-associated promoter-adjacent DNA hypermethylation at *TBX15* fine-tunes expression of this gene by downmodulation. Our findings have implications for better understanding differentiation within the SkM lineage and the transcriptional and epigenetic changes in SkM that occur with exercise and aging [20].

## 2. Results

### 2.1. Myoblast-associated DNA hypermethylation around an unmethylated promoter region was positively associated with expression of TBX15, TBX18, TBX2, TBX3, and TBX1

RNA-seq databases show that human *TBX15* is preferentially transcribed in SkM tissue (highest expression), myoblasts, myocytes, fibroblast-type cells in SkM, as well as in some other cell types in postnatal tissues (smooth muscle cells and adipocytes; Figures 1A and S1; Table S1). Moreover, among fetal tissues, SkM shows the highest expression of *TBX15* (Table S2). However, there are considerable differences in transcript levels for *TBX15* in human SkM depending on the anatomical origin of the tissue (Table S2). *TBX15*’s closest related T-box encoding gene is *TBX18* (Figure 1B) [3], which has less of a preference for expression in myoblasts than does *TBX15* (Table S3).

**Figure 1.**
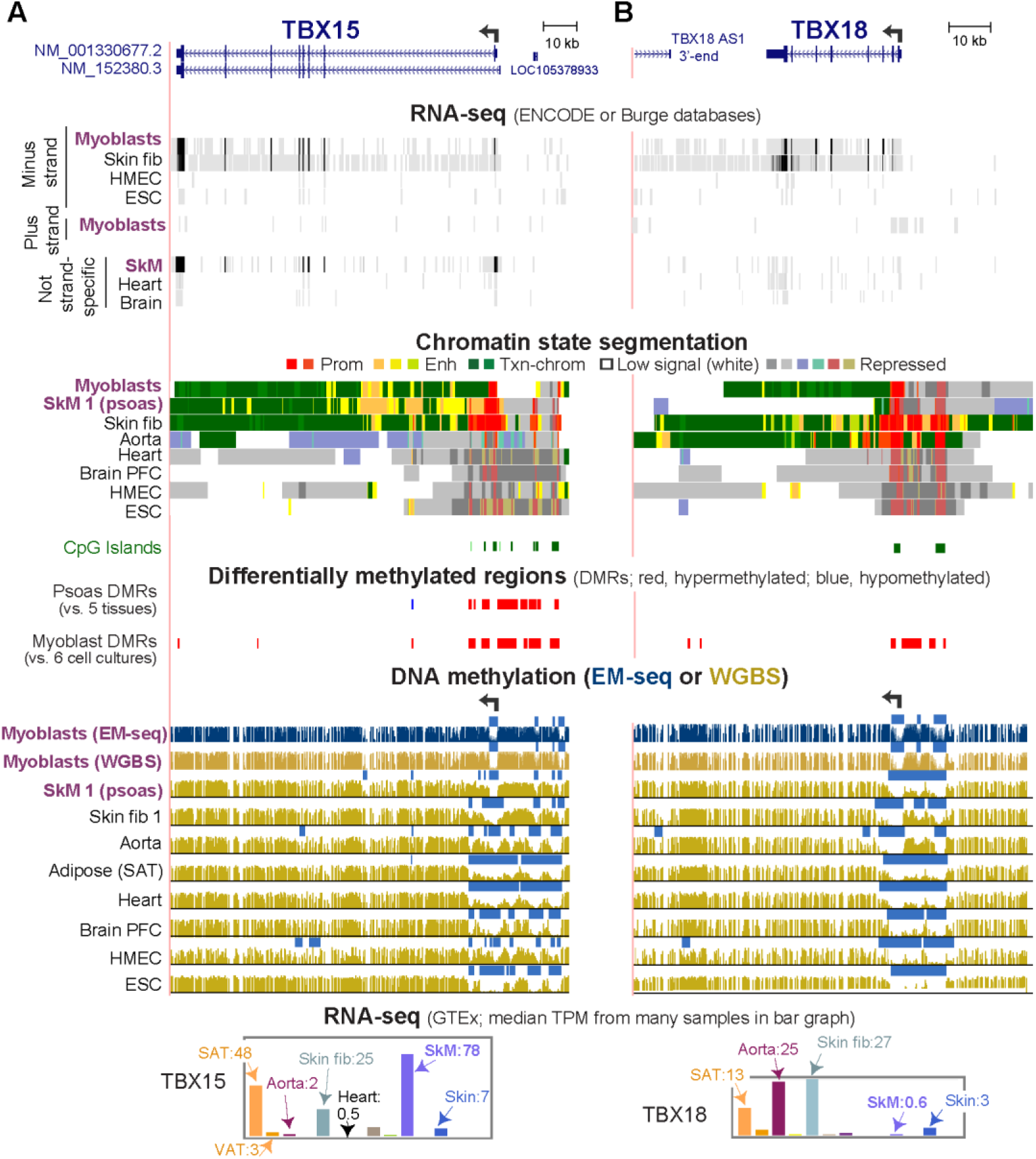
DNA hypermethylation in myoblasts borders the active, unmethylated promoter regions of *TBX15* and *TBX18*. (**A**) *TBX15* (chr1:119,423,765-119,554,608) and (**B**) *TBX18* RefSeq gene structures (chr6:85,410,477-85,505,693). Strand-specific RNA-seq for myoblasts, foreskin fibroblasts (Fib, Fib 3), mammary epithelial cells (HMEC), and H1 embryonic stem cells (ESC) and not strand-specific RNA-seq for skeletal muscle (SkM), heart and brain. Color-coded chromatin state segmentation indicates promoter or mixed promoter/enhancer (prom), enhancer (enh), or repressed types of chromatin, actively transcribed chromatin (txn chrom), or chromatin with little or no signal for H3K27ac, H3K27me3, and H3K4/H3K9 methylation. Significant tissue-specific (SkM, psoas) DMRs and cultured cell-type specific (myoblast) DMRs are shown. Methylome profiles are depicted in gold for WGBS and, in dark blue, for EM-seq; regions having significantly lower methylation relative to the same genome [21] are shown by light blue bars. Myoblast 3 cell strain was used for EM-seq and WGBS and Skin fib 1 (foreskin fibroblasts) for WGBS. GTEx RNA-seq expression profiles are displayed as linear-scale TPM bar graphs with some of the median TPM values from biological replicates indicated. SAT, subcutaneous adipose; VAT, visceral adipose; heart, left ventricle. All tracks are from the UCSC Genome Browser (hg19) except for the GTEx bar graphs and are aligned.

In myoblasts and SkM, expression of *TBX15* was positively correlated with DNA hypermethylation both immediately upstream and downstream of the unmethylated promoter region (Figure 1A). *TBX18* displayed a similar correlation but only for myoblasts (Figure 1B). In addition, both genes in skin fibroblasts showed a positive association between transcription and DNA hypermethylation upstream of the promoter although the hypermethylation was not as strong as in myoblasts. *TBX18* exhibited this association for aorta too. As expected, a lack of methylation at the TSS does not suffice for appreciable T-box gene expression, as seen for *TBX15* in heart and *TBX18* in SkM (Figure 1B). In tissues not expressing these two genes, the promoter regions, which contained mostly unmethylated DNA, exhibited repressive histone H3 lysine-27 trimethylation (H3K27me3). For our DNA methylation analyses, DMRs were identified by comparing methylomes from SkM (psoas) with five diverse tissues [22] and by comparing three myoblast cell strains with six types of non-cancerous cell cultures (Figure S2). Methylomes for the non-myoblast cultures and for all the tissues had been determined by wholegenome bisulfite sequencing (WGBS) [23]. The myoblast methylomes were generated by the recent enzymatic methyl-seq methodology (EM-seq) [24]. One myoblast cell strain (Myoblast 3) was analyzed by WGBS as well as EM-seq. Both methods gave similar DNA methylation profiles (Figure 1A, gold *vs*. blue methylome tracks for the Myoblast 3 cell strain). Cis-acting transcription-control chromatin elements, such as, repressive chromatin (H3K27me3 or H3K9me3), active promoters (H3K27 acetylation, H3K27ac, plus H3K4me3) and active enhancers (H3K27ac plus H3K4me1) had been inferred from chromatin segmentation state profiles derived from whole-genome maps of diagnostic histone modifications (Roadmap Project [19]).

All 17 members of the T-box family of genes are expressed postnatally in highly cell- or tissue-specific patterns (Table S1) consistent with their important roles in development [3,25]. Like *TBX15* and *TBX18*, three other T-box genes, *TBX2, TBX3*, and *TBX1*, were more highly expressed in myoblasts than most or all the five other diverse cell culture types in an ENCODE database (Table S3). In myoblasts, *TBX2*, and *TBX3* also displayed *TBX15*-like DNA hypermethylation bordering both sides of a constitutively unmethylated promoter region (Figures 1, S3, and S4). This promoter-adjacent DNA hypermethylation was not seen in non-expressing cell cultures or tissues nor was it observed in lung fibroblasts, which had by far the highest levels of expression of the six examined cell cultures and enhancer or promoter-type chromatin covering the whole gene, promoter, and promoter-upstream region. At *TBX1*, a myoblast-associated hypermethylated DMR (Myob-hyperm DMR) was adjacent to the upstream border of its unmethylated proximal promoter region (Figure S3). There was high methylation immediately adjacent to this promoter’s downstream border but this methylated region was not a DMR because it was present in most examined cell types. About 1 kb downstream from the promoter, a region of myoblast-associated hypermethylation (another Myob-hyperm DMR) was seen. Although there are RefSeq isoforms of *TBX1* with an alternate distal promoter, examination of RNA-seq databases at the UCSC Genome Browser having many diverse samples provides evidence for use of only the proximal promoter (Figures S3A and data not shown).

*TBX20, TBX4*, and *TBX5* were among the T-box genes not expressed in myoblasts (Table S3). This transcription silencing in myoblasts correlated with Myob-hyperm DMRs covering their promoter regions instead of being only adjacent to them (dotted boxes, Figures S4 and S5). These three genes also exhibited repressive H3K27me3 at the promoter region. CpG islands (CGI) are present in the promoter regions of the five myoblast-expressed *TBX* genes as well as the three above-mentioned myoblast-repressed (Figures 1, 2, and S3-S5). In the myoblast-repressed genes, the promoter/CGI-overlapping DMRs could contribute to the gene repression. As reported for some intragenic CGIs [26], the CGI-overlapping Myob-hyperm DMRs in these T-box genes in various non-expressing cell types often overlapped bivalent (mixed H3K27me3-repressed and H3K4methylated-enhancer/promoter-type) chromatin instead of only H3K27me3 chromatin (Figures 1, S3-S5) reflecting their potential for promoter or enhancer activity. We focused the rest of the study on the 5’ end of *TBX15*, and its strong promoter-adjacent (but not overlapping) DNA hypermethylation that was found only in expressing cells.

**Figure 2.**
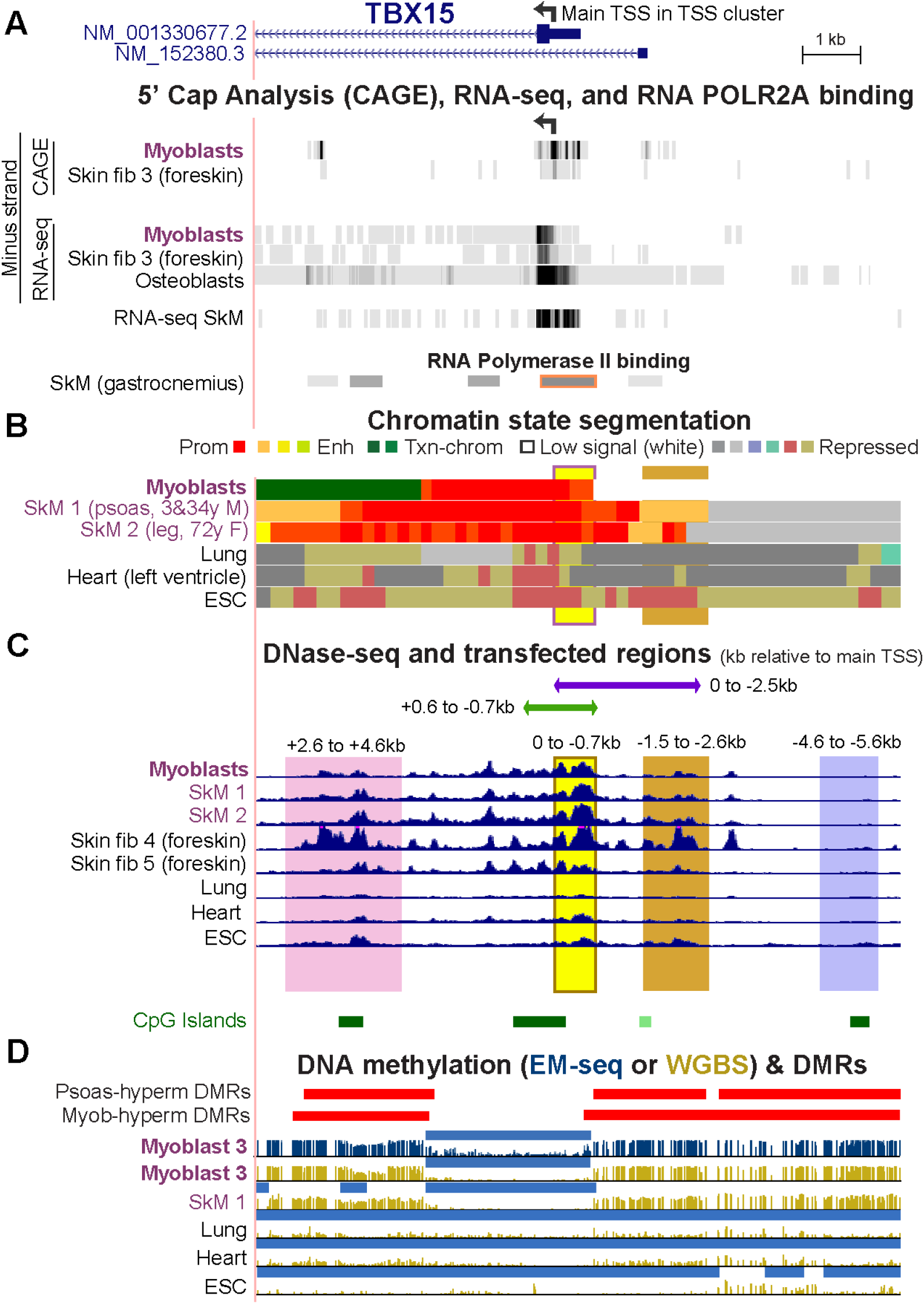
The myoblast-hypermethylated DMRs adjacent to the unmethylated *TBX15* promoter region overlap psoas-hypermethylated DMRs. (**A**) The 5’ end of *TBX15* (chr1:119,525,320-119,536,521) showing the minus-strand signal from 5’ cap analysis of gene expression (CAGE) and/or RNA-seq of myoblasts, foreskin fibroblasts, and osteoblasts; not strand-specific RNA-seq for SkM of unknown body location (SkM 9); and the gastrocnemius muscle ChIP-seq signal for the large subunit of RNA polymerase II (ENCODE 3). (**B**) Chromatin state segmentation as in Figure 1. (**C**) DNaseI hypersensitivity profiles and the six cloned regions for transfection assays shown by horizontal bars or vertical shading. (**D**) Hypermethylated DMRs for SkM (psoas) *vs*. five heterologous tissues and for myoblasts *vs*. six varied cell cultures as well as methylome profiles and low methylated regions (light blue bars) as in Figure 1.

### 2.2. DNA sequences that were part of myoblast hypermethylated DMRs near the TBX15 promoter region display promoter or enhancer activity upon transfection into myoblasts

To understand the function of cell-type specific DNA hypermethylation around core unmethylated promoter regions, we tested the transcription regulatory activity of *TBX15* DNA sequences from Myob-hyperm DMRs in transient transfection assays using reporter gene constructs (Vector 1 or 2) for transfection into C2C12 myoblasts (Figures 2 and 3). None of the cloned sequences overlapped interspersed DNA repeats [23]. First, we ascertained the approximate location of the main *TBX15* TSS in myoblasts and skin fibroblasts so that we could use that site as a reference point for the cloned sequences. The 5’ ends of the two RefSeq isoforms (RefSeq Curated, 2022) are separated by 1.2 kb (Figure 2A). However, according to cap analysis of gene expression (CAGE) and/or RNA-seq profiles from the ENCODE or RoadMap Projects, neither of these 5’ ends is the best description of the *TBX15* TSS in myoblasts, skin fibroblasts, osteoblasts (which also preferentially express this gene, Table S3), and several types of SkM (Figure 2A and data not shown for SkM RNA-seq at the UCSC Genome Browser [19,23]). We used the center of the strongest CAGE signal for myoblasts and skin fibroblasts, chr1:119530511 (hg19), as the nominal TSS. This site (broken arrow in Figures 1A and 2A), which we refer to as the main TSS, is inside an RNA Pol II binding site found in SkM (Figure 2A, POLR2A large subunit binding) and the unmethylated 5’ region of *TBX15* in myoblasts and SkM (Figure 2D). This TSS and the 0.5-kb upstream TSS of isoform NM_0011330677 are predicted to encode the same main TBX15 protein of 602 amino acids that is observed in normal skin fibroblasts [27]. The cluster of CAGE-determined 5’ ends of myoblast, skin fibroblast, and osteoblast transcripts is in promoter chromatin and adjacent to a DNaseI hypersensitive site and overlaps a CGI (Figure 2A-C).

**Figure 3.**
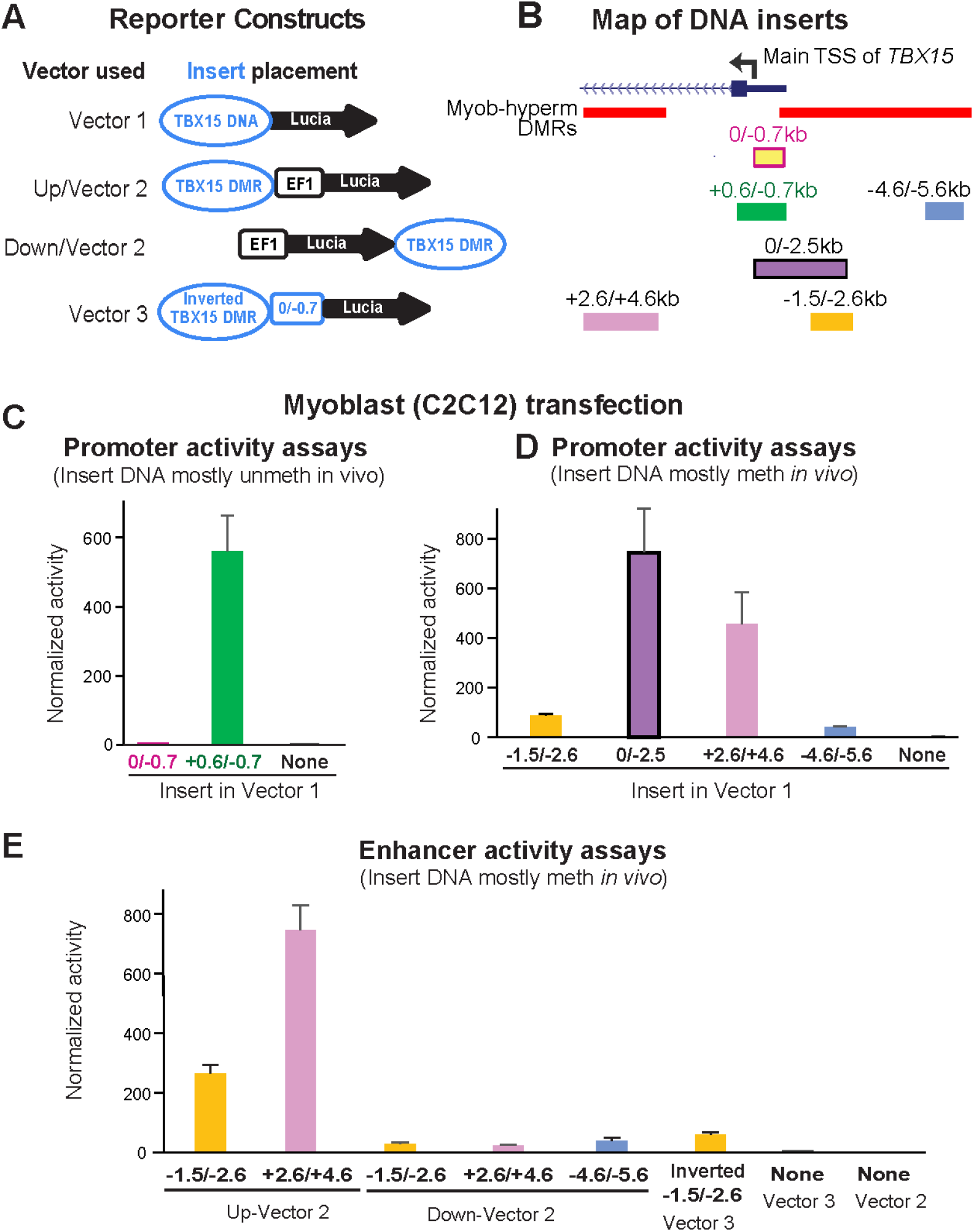
Myoblast-hypermethylated DNA sequences adjacent to the unmethylated promoter displayed strong promoter or enhancer activity from reporter constructs transfected into myoblasts. (**A**) The constructs used for transfection are diagrammed. Vector 1 is the promoter-less pCpGfree-Lucia plasmid; Vector 2 is the analogous plasmid with a modified human EF-1α-derived minimal promoter (EF1). *TBX15* DNA, the constitutively unmethylated promoter region or DMR sequences adjacent to it. (**B**) Endogenous location of the *TBX15* inserts used in reporter gene assays (Figure 2). Myob-hyperm DMRs are named according to the beginning and end positions relative to the main TSS of *TBX15*. (**C, D,** and **E**) Reference plasmid-normalized luciferase activity from C2C12 myoblasts transiently transfected with plasmids containing inserts shown in Panel **B** in assays for promoter (Panel **C** and **D**) or enhancer (Panel **E**) activity. In human myoblasts, the inserts were mostly unmethylated (Panel **C**) or highly and specifically methylated (Panels **D** and **E**) *in vivo* but became unmethylated upon cloning. Results from transfections are the averages from at least three independent experiments; error bars for standard error. Meth, methylated; unmeth, unmethylated.

We first tested promoter activity using a CpG-free promoter-less Lucia vector (Vector 1) and inserts from constitutively unmethylated promoter region sequences from the main *TBX15* TSS to 0.7 kb upstream (0/−0.7 insert; Figure 3A and B). Upon transient transfection into C2C12 myoblasts, luciferase reporter activity was almost undetectable from this plasmid and was not significantly greater than the background luciferase activity from Vector 1 (Figure 3C). Although the 0/−0.7 plasmid lacked promoter activity in transfected myoblasts, the endogenous sequences in human myoblasts and in SkM display tissue-specific promoter chromatin and DNaseI hypersensitivity (yellow highlighting, Figure 2B and C). Secondly, we enlarged the test insert by 0.6 kb of constitutively unmethylated sequences using a +0.6/−0.7 insert instead of the 0/−0.7 insert. With this construct, strong promoter activity was seen in the transfected myoblasts (Figure 3C). These findings suggest that endogenous sequences from the TSS to 0.7 kb upstream in myoblasts and in SkM participate in promoter activity through cooperation with neighboring, TSS-downstream unmethylated DNA sequences that lack their own promoter activity. This explanation is supported by the finding of more promoter chromatin and unmethylated DNA immediately downstream of the TSS than upstream of the TSS (Figure 2B and D).

Next, we determined the promoter activity of an enlarged TSS-upstream region from the main TSS to −2.5 kb (0/−2.5; Figure 3B). This plasmid displayed high luciferase activity in myoblast transfectants (Figure 3D) even though its constituent DNA sequences are the promoter-inactive 0/−0.7 sequence and 1.8 kb of *TBX15*-upstream DNA, a sequence that is highly methylated in endogenous human myoblast DNA and has no overlapping promoter chromatin and only low DNaseI hypersensitivity (brown highlighting, Figure 2B and C). Interestingly, quantitation of the WGBS profiles in this region (−0.7 to −2.5 kb, chr1:119,531,244-119,533,033, hg19) showed that the psoas SkM sample had significantly less methylation (p = 2E-27) than the Myoblast-3 cell strain (average methylation 91% for myoblasts, based on our WGBS data, and 67% for SkM, using Roadmap WGBS data [19]). Accordingly, SkM (psoas) tissue displayed specific acquisition of enhancer or promoter chromatin in this region not seen in myoblasts (Figure 2B).

Both a 1.1-kb insert that came only from a Myob-hyperm DMR (−1.5/2.6; Figure 3B) and a far-upstream 1.0-kb insert from the same DMR (−4.6/5.6) displayed promoter activity in transfected myoblasts relative to the background activity of Vector 1 alone (T-test for each construct *vs*. vector only, p < 1E-10). However, the promoter activity from the −1.5/−2.6 insert was much less than from the larger 0/−2.5 insert (Figure 3D). This again indicates the ability of the 0/−0.7 DNA sequence to cooperate with adjacent *TBX15* promoter region sequences to confer promoter activity, even though the 0/0.7 sequence had no promoter activity by itself in transfected reporter constructs.

When the −1.5/−2.6 DNA sequence was assayed for enhancer activity by inserting it upstream of a minimal EF1 promoter in Vector 2 (Figure 3A and B), high luciferase activity was seen in myoblast transfectants (Figure 3E). The strongest enhancer activity was observed for a 2-kb Myob-hyperm DMR from *TBX15* intron 1 (+2.6/+4.6, Figure 3E). The genomic location of this DMR is at the downstream border of the constitutively unmethylated promoter region (Figure 2C and D). Usually, inserts tested for enhancer activity in reporter gene constructs are inserted upstream of a minimal promoter in a reporter plasmid, as was done in the experiments demonstrating high enhancer activity for the +2.6/+4.6 region (+2.6/+4.6 Up-Vector2; Figure 3E). When enhancer activity was tested more rigorously in constructs containing the insert downstream of the reporter gene, significantly enhanced reporter gene activity was observed (+2.6/+4.6 Down-Vector2, Figure 3E; T-test *vs*. Vector 2, p = 4E-5) but much less than that from the promoter-upstream insertion. The −1.5/−2.6 and −4.6/−5.6 inserts also gave low, but significant, enhancer activity when tested downstream of the reporter gene (Figure 3E; T-test *vs*. Vector 2, p =1E-4 and 0.01, respectively). We previously studied the activity of the *MYOD1* core enhancer by cloning it downstream of the reporter gene in the same CpG-free, minimal EF1 promoter vector and obtained strong activity from comparable transfection assays [28], thus indicating no technical problem with the use of the downstream cloning site in this vector. These results suggest context-dependent upregulation by *TBX15* Myob-hyperm DMR sequences when tested as unmethylated sequences in reporter constructs.

### 2.3. Much lower enhancer and promoter activity was seen for transfected TBX15 TSS-upstream or downstream sequences in non-myoblast vs. myoblast host cells

The above-described reporter gene constructs containing *TBX15* promoter-upstream or downstream sequences were also transfected into MCF-7 cells, a breast cancer-derived epithelial cell line. As expected from the widespread use of the MCF-7 cell line for transfection assays, these cells were highly transfectable, which we verified by testing promoter activity of DNA from the broadly expressed *IRS1* promoter region (data not shown). However, when reporter constructs containing Myob-hyperm DMR-derived inserts were used for transfection of MCF-7 cells, the transfectants had much lower reporter gene activity than that of analogous C2C12 myoblast transfectants (Figure 4A and B *vs*. Figure 3D and E, note different scales). The +2.6/+4.6 Myob-hyperm DMR had 109- and 69-fold lower transcription-promoting activity in transfected MCF-7 cells than in transfected myoblasts in promoter and enhancer test assays, respectively. Comparable decreases for the −1.5/−2.6 Myob-hyperm DMR sequences were 10-to 12-fold lower in MCF-7 than in myoblast transfectants.

**Figure 4.**
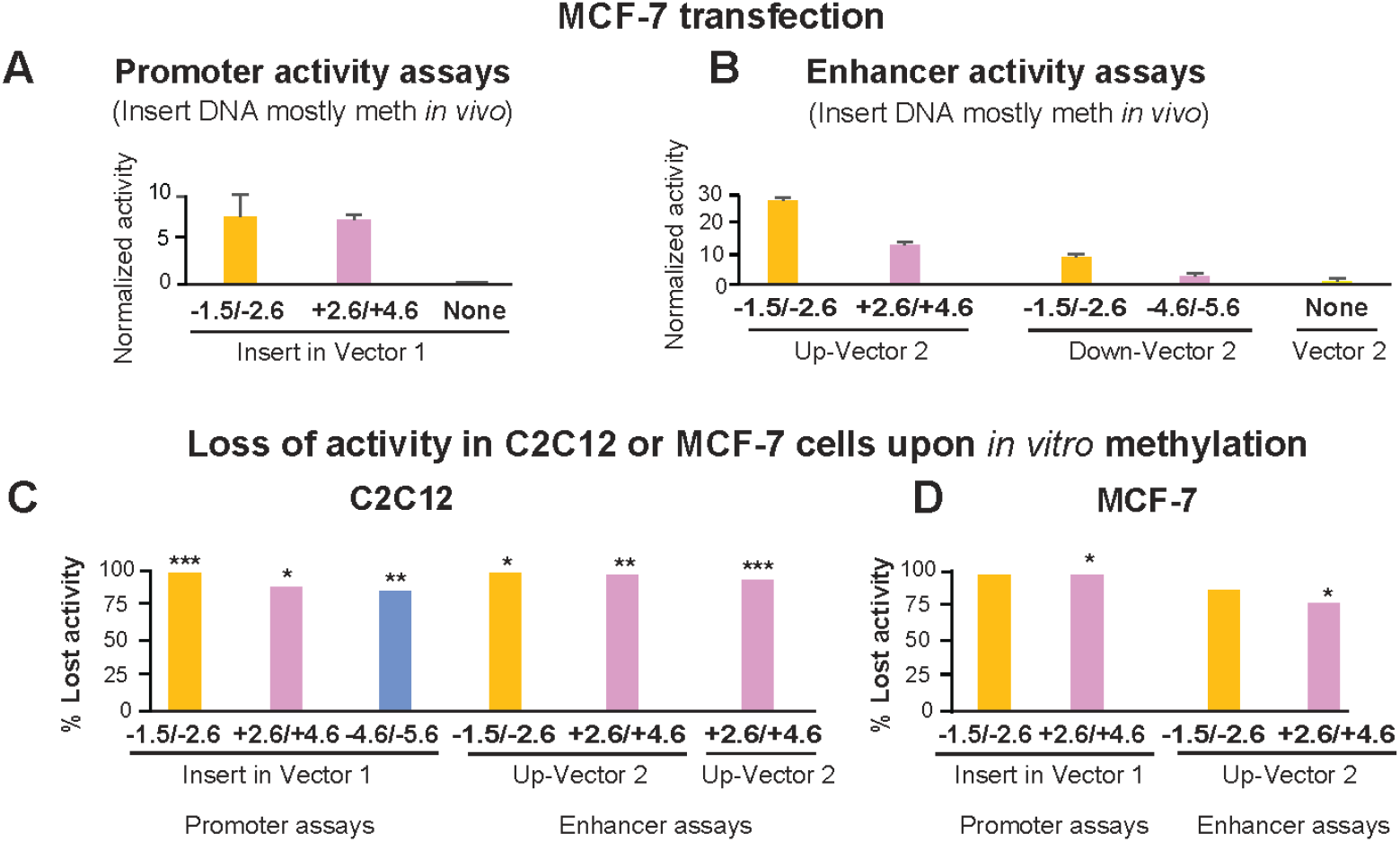
The promoter or enhancer activity of DMRs from the 5’ end of *TBX15* was strongly decreased by targeted *in vitro* methylation or transfection into MCF-7 cells instead of myoblasts. Transfection was done with promoter-assay (**A**) or enhancer-assay (**B**) constructs as in Figure 3 but with the MCF-7 breast carcinoma cell line instead of C2C12 myoblasts as the host cells. Note the very different scales for normalized luciferase activity in Figures 3 *vs*. 4. The normalized luciferase activity lost in C2C12 myoblasts (**C**) or MCF-7 cells (**D**) transfected with *in vitro* methylated compared with mock-methylated reporter gene constructs. The difference between methylated and unmethylated DNA was significant at p <0.05 (*), p <0.01 (**) or p <0.001 (***). *In vitro* CpG methylation was targeted only to the insert.

### 2.4. In vivo-like methylation by M.SssI CpG methyltransferase decreased enhancer or promoter activity of TBX15 DMRs upon transfection

The reporter gene constructs containing inserts that were highly methylated in human myoblasts were assayed for the effects of CpG methylation on their promoter or enhancer activity. In the above-described experiments, the myoblast CpG methylation is lost upon cloning in *E. coli*. Because the reporter gene vectors that we used had been engineered to contain no CpGs, M.SssI-catalyzed methylation (which is CpG-specific) could only occur in the inserts; therefore, there could be no effects on reporter gene expression from methylation of the reporter gene or of the rest of the vector [29].

The −1.5/−2.6 DMR sequence-containing constructs for testing promoter activity or enhancer activity lost 97 to 100% of their activity upon M.SssI methylation relative to mock-methylated controls in transfected myoblasts (Figure 4C). Analogous assays for promoter or enhancer activity of +2.6/+4.6 or −4.6/−5.6 DMR inserts in transiently transfected C2C12 cells gave losses of activity of 86 – 89% (Figure 4C). Methylation *in vitro* to resemble the high extent of CpG methylation of these endogenous sequences in human myoblasts also resulted in the loss of most activity of these constructs when they were transfected into MCF-7 cells (Figure 4D). In summary, the promoter activity seen for all the DMR sequences tested was largely or completely dependent on these sequences not being highly methylated, as they are in human myoblasts

### 2.5. Variable epigenetics at the 5’ end of TBX15 in some skeletal muscle samples, myoblast and skin fibroblast cell strains, adipocytes, and cancer cell lines

We looked at the epigenetics *in vivo* of the cloned Myob-hyperm DNA regions in additional samples of myoblast and skin fibroblast cell strains, SkM tissue, and in cancer cell lines. We used our previous DNA methylation profiles from RRBS [30] of myoblasts supplemented with other ENCODE RRBS profiles [31]. RRBS only detects the methylation state of 5% or less of CpGs genome-wide [32] but regions of high CpG density, like the 5’ end of *TBX15*, are overrepresented (Figure 5A and B) in RRBS profiles. Myoblast 3 and Myoblast 7 displayed similar methylation patterns in RRBS profiles to those seen in WGBS or EM-seq profiles of Myoblast 1, 3, and 6 (Figures 2D and 5C). In contrast, RRBS revealed that Myoblast 8, SkM 7, and SkM 8 lacked most of the high DNA methylation observed in the other myoblast and SkM samples around the constitutively unmethylated *TBX15* promoter core (Figures 5B, C and 2D). Myotubes, which we obtained by *in vitro* differentiation of the myoblast cell strains, shared indistinguishable *TBX15* DNA hypermethylation profiles (Figure 5B) although the myotubes had 1.8 times the RNA levels as myoblasts (RNA-seq data, not shown).

**Figure 5.**
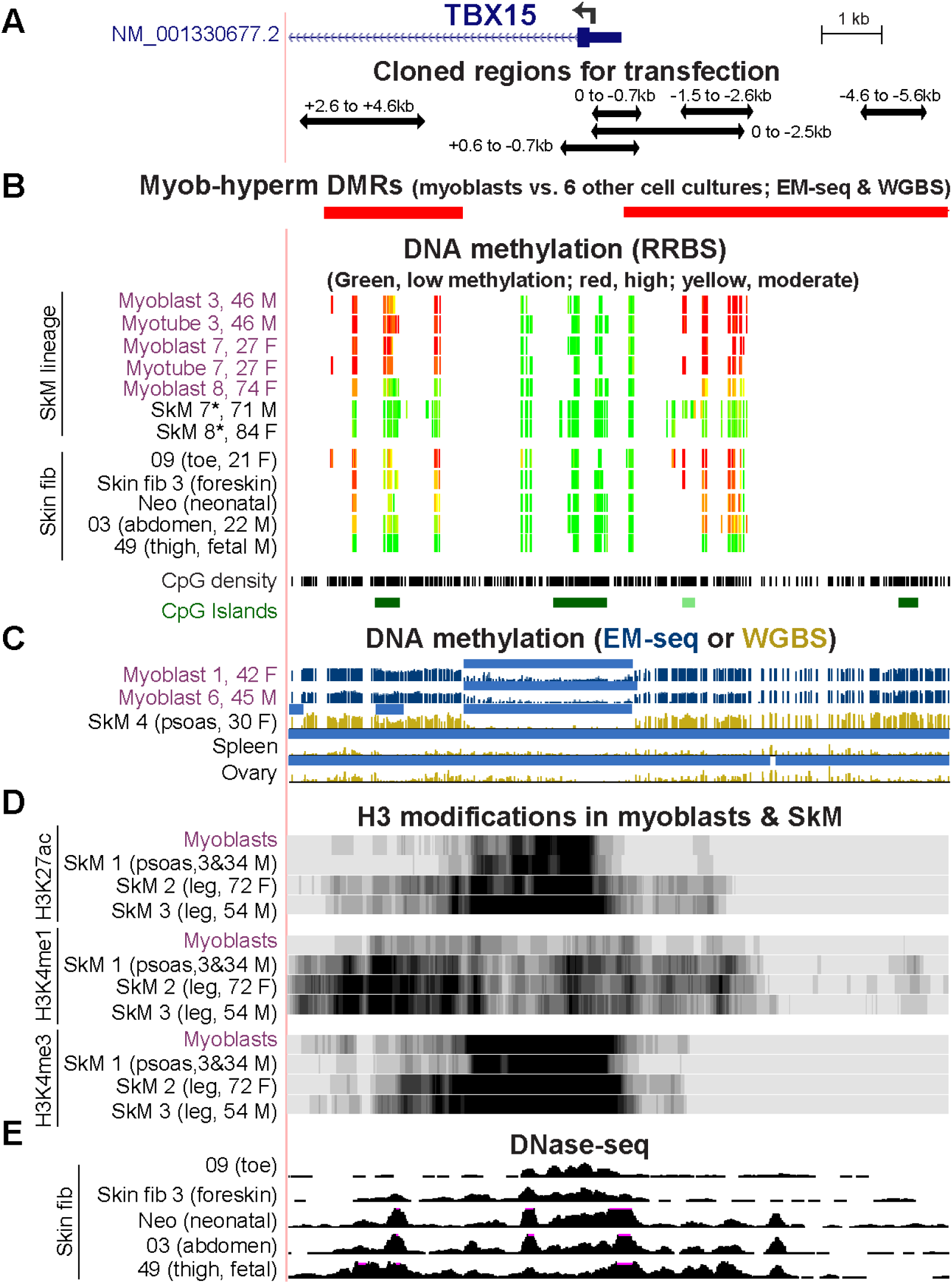
Some sample-specific differences in epigenetic profiles at the 5’ end of *TBX15*. (**A**) Positions of cloned regions used for transfection (chr1:119,525,320-119,536,521; only the RefSeq Select isoform is shown). (**B**) WGBS- and EM-seq determined DMRs, RRBS profiles of DNA methylation, CpG density, and CpG islands. (**C**) EM-seq and WGBS profiles of DNA methylation with light blue bars denoting low methylated regions as in Figure 1. (**D**) H3K27ac, H3K4me1, and H3K4me3 signal and (**E**) DNaseI hypersensitivity for the indicated samples with vertical viewing ranges of 0-20, 0-10, 0-20, and 0-30, respectively. The ages and genders of donors of myoblast, SkM, and skin fibroblast samples are given, where known; Neo, neonatal skin fibroblasts were not further identified [23]; *, SkM from an unspecified body region.

The SkM tissue samples that exhibited myoblast-like DNA hypermethylation around the 5’ end of *TBX15* were from young donors (SkM 1, a mixture of 3-y M and 34-y M) while the samples lacking most of this hypermethylation (Figure S2) were derived from elderly individuals (71- to 84-y M or F; unknown type of SkM). Another complicating factor is that Myoblast 8 was derived from a 74-y patient with inclusion-body myositis so that the disease state might have influenced *TBX15* DNA methylation. In RRBS methylomes, myoblasts and myotubes from one of two young patients with facioscapulohumeral muscular dystrophy displayed reduced amounts of methylation at the Myob-hyperm DMRs while the other sample exhibited the kind of hypermethylation seen in most myoblast and myotube samples (Figure S6). Although the RRBS profiles at the 5’ end of *TBX15* reveal the Myob-hyperm DMRs (Figure 5), the WGBS and EM-seq profiles show the specificity of these DMRs for myoblasts and skin fibroblasts more clearly than do RRBS profiles (Figures S2 and S6).

WGBS methylomes were publicly available for only two SkM samples, both of which were from psoas muscle (SkM 4, 30-y F and SkM 1) [19]. These methylomes were similar in the vicinity of *TBX15* (Figures 2D *vs*. 5C). From available histone methylation and chromatin state and histone H3 modification profiles for the psoas SkM 1, leg muscle SkM 2 (72-y F) and leg muscle SkM 3 (54-y M), there was more H3K27ac (indicative of active promoter or enhancer chromatin) in the cloned DMRs in the leg muscle samples than in the psoas or myoblast samples (Figure 5D). The H3K27ac profiles of SkM and myoblasts indicate the presence of a super-enhancer, a large cluster of enhancer and/or promoter chromatin regions [33]. It spanned ~32 and 46 kb from the promoter region through much of the large intron 1 for psoas and leg muscle, respectively (Figures 1 and S6). This superenhancer was not seen in myoblasts [34].

Although the small number of available SkM samples analyzed for their epigenetics at the 5’ end of *TBX15* exhibited no correlation with gender, RNA-seq analysis of 543 male and 260 female gastrocnemius muscle samples (GTEx project [35]) showed that the median TPM (transcripts per million) value for females was 16% higher than that for males (rectangle, Figure S7). A substantial gender-difference in TPM levels was also noted for *TBX1* in this database, but not for the *TBX15* neighbor *WARS2*. The statistical and biological significance of this finding is uncertain.

Like myoblasts, almost all the primary skin fibroblast cell cultures displayed DNA hypermethylation immediately upstream and downstream of an unmethylated *TBX15* TSS-overlapping region as analyzed by RRBS or WGBS [15,36] (Figures 1A and 5B and S2). These skin fibroblast cultures were derived from various body depots (Figure 5B). Like the myoblast primary cultures, primary cultures of skin fibroblasts (from postnatal leg, temple, scalp, breast, abdomen, back, and neonatal foreskin dermis) express *TBX15* at moderate levels (Figure 1A, top and bottom panels, and data not shown from ENCODE and Roadmap databases). Skin fibroblasts varied in their extent of DNA hypermethylation in the TSS −20 to +9 kb region (Figure 5B) even among biological replicates of foreskin fibroblast primary cultures (Figures 1A and S2B). The one examined skin fibroblast cell strain (Skin fib 49) that was fetal in origin was exceptional in displaying no hypermethylation in this region. Among the skin fibroblast cultures, there was more open chromatin (DNaseI hypersensitivity) in the regions of less DNA methylation (Figure 5B and E, *e.g*., toe and foreskin *vs*. fetal thigh).

Another cell type with highly varied epigenetics according to the subtype is uncultured adipocytes. As previously reported in a WGBS and RNA-seq study of adipocytes [37], methylation at *TBX15’s* 5’ end differed strongly between adipocytes from the same individual derived from either from subcutaneous adipose tissue, which highly expresses *TBX15*, or from visceral adipose tissue, which shows only low levels of *TBX15*. Promoter border DNA hypermethylation was associated with the more highly expressing adipocytes. We found these subcutaneous adipose hypermethylated DMRs overlapped the Myob-hyperm DMRs adjacent to the constitutively unmethylated *TBX15* promoter (Figure S2B).

Myob-hyperm DMRs were located not only immediately around the unmethylated *TBX15* promoter region but also as part of an extended cluster of DMRs in myoblasts, psoas muscle, and a skin fibroblast cell strain (Figures 1A and S2B). Another prominent SkM-lineage related difference in *TBX15* epigenetics is that there was additional cell type-specific enhancer or weak enhancer chromatin within the gene body or far upstream or downstream of its promoter that was associated with *TBX15* tissue expression profiles (Figures 1, S2, S6, and S8). This included enhancer chromatin overlapping SkM-specific DNA hypomethylation in the large gene desert downstream of the gene as far as 0.5 Mb distant from the gene in both psoas and leg muscle (Figure S8A).

Many cancer cell lines that did not express *TBX15* differed from normal cell strains in having high levels of DNA methylation throughout the region from TSS −20 to +9 kb, as seen in RRBS profiles [31] (Figure S6). However, other cancer cell lines without this promtoer-overlapping DNA hypermethylation still did not express *TBX15* [38], like most non-transformed cell strains. We found RNA-seq [38] and methylome data [31] for one cancer cell line that expressed *TBX15*. Importantly, this cell line, U87 astrocytoma cells, was the only cancer cell line exhibiting *TBX15*-like DNA hypermethylation around an unmethylated TSS region (Figure S6) with the exception of HepG2 cells. HepG2 cells do not initiate transcription at the canonical 5’ end of *TBX15* and instead use a liver promoter located in exon 6 of the main isoform (Figures S6A, green signal, and S8A; CAGE and GTEx data not shown [23,35]). Interestingly, like tissues which do not express *TBX15* from any promoter, liver lacks DNA hypermethylation at or around the main promoter region (Figure S6E).

### 2.6. Transcription factor binding sites in the 5’ TBX15 region help explain enhancer and promoter activity observed in transfection experiments and chromatin epigenetic profiles

Because there are only a very small number of genome-wide profiles of TF-directed chromatin immunoprecipitation-next gen sequencing (ChIP-seq) from myoblasts (mostly POLR2A, MYOD, MYF5, CTCF), we looked for evidence of TF binding at the above-described cloned *TBX15* regions in various cell types and examined predicted transcription factor binding sites (TFBS) in these regions. From the Unibind human TFBS database, which is based upon TF ChIP-seq profiles combined with TFBS predictions [39], we found binding sites in the cloned regions at the 5’ end of *TBX15* for MYOD, one of the four TFs found specifically in the SkM lineage. One of the MYOD sites in myoblasts and rhabdomyosarcoma cells (cancers derived from myogenic progenitor cells) is within the *TBX15* +2.6/+4.6 Myob-hyperm DMR (Figure 6A and B; Table S4). MYOD also occupied two sites in an adjacent unmethylated region in myoblasts, one of which overlapped a large DNaseI-hypersensitive peak in myoblasts. All three MYOD sites also overlapped POLR2A binding subregions in gastrocnemius SkM (Figures 2A and 6B). The two clustered MYOD sites bind to homologous mouse DNA sequences as determined from C2C12 myoblast Myod ChIP-seq profiling [28,40]. Importantly, the mouse dataset gives the relative amount of binding of MYOD. Both C2C12 Myod binding sites in *TBX15* intron 1 were only weak sites with binding scores of 13 and 19 compared to 91 – 168 for strong Myod sites in enhancer chromatin far upstream of the *Myod1* gene in the same ChIP-seq profile [40].

**Figure 6.**
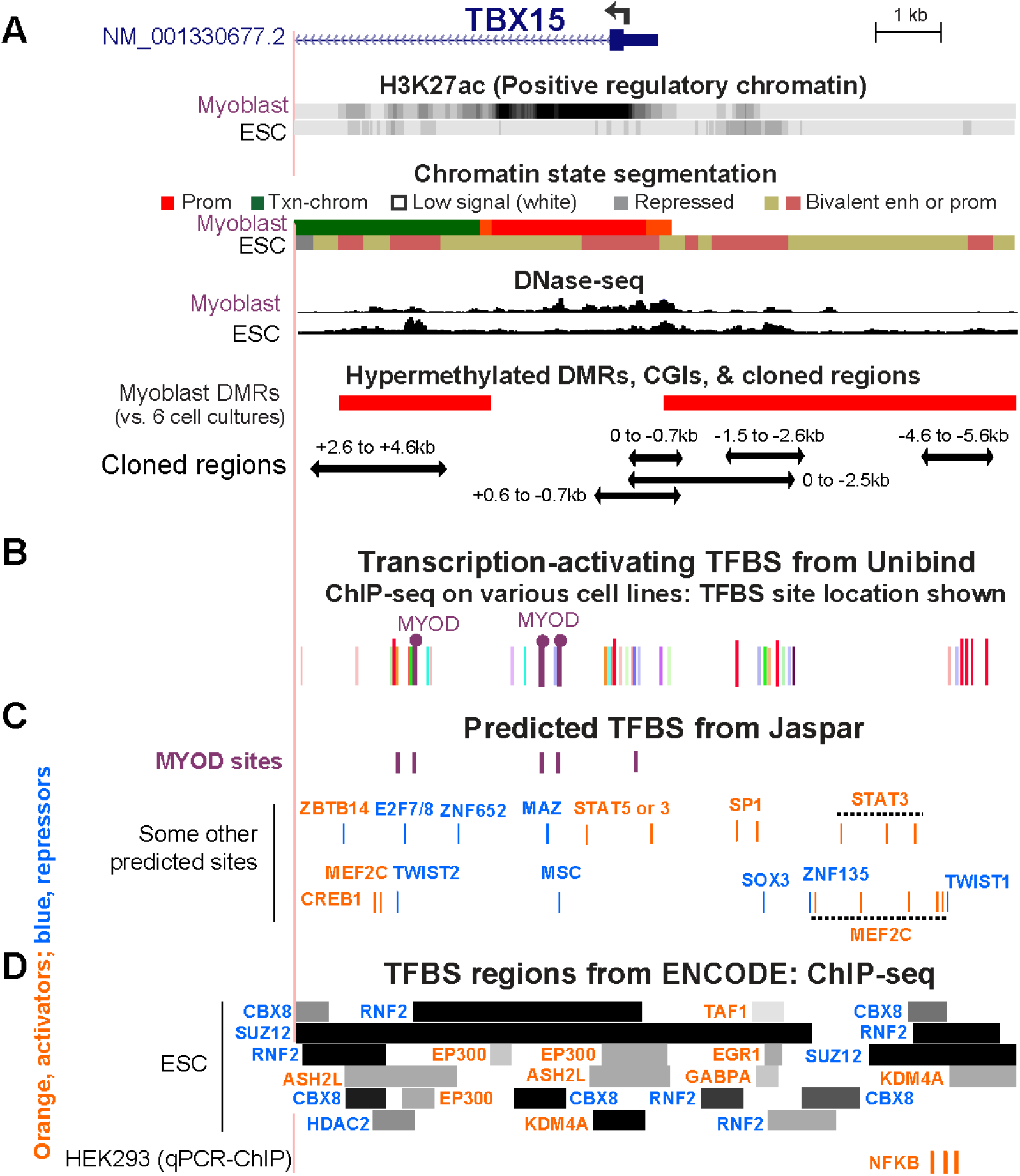
Overlap of cloned regions from the 5’ end of *TBX15* with transcription factor binding sites. (**A**) Myoblast and embryonic stem cell (ESC) chromatin epigenetics, Myob-hyperm DMRs, and cloned regions used for reporter gene assays as in Figure 2 (chr1:119,525,320-119,536,521). (**B**) Bars, the locations of representative TFBS from combined ChIP-seq data and TFBS predictions (Unibind database; see Table S4 for details). Binding sites found by ChIP-seq for MYOD, a myogenesis-associated TF, in myoblasts or rhabdomyosarcoma cells are shown by lollipops. (**C**) Predicted TFBS for MYOD (JASPAR database). (**D**) ChIP-seq-determined regions of binding of TFBS to ESC (ENCODE 3) and sites of experimentally determined binding of NFKB to HEK293 cells [41] are shown. Binding regions indicated by dark colored segments denote strong TF binding. In Panels **C** and **D**, TF labels for transcription-activating TFs are shown in orange and for repressing TFs [8] in blue.

Binding profiles are available for many cell types for the CCCTC-Binding Factor (CTCF), a protein that mediates chromatin looping as well as being a sequence-specific TF. A strong constitutive CTCF binding site (Figure S2C) was seen immediately upstream of the cluster of Myob/SkM-hyperm DMRs between the 5’ end of *TBX15* and the 3’ end of its neighbor *WARS2*, a broadly expressed gene (Table S3B). A weak CTCF site that was highly cell type-specific was seen towards the 3’ end of *TBX15* in 1 in myoblasts, myotubes, skin fibroblasts, and osteoblasts, all of which express *TBX15* (Figure S2). Weaker CTCF sites were observed in myoblasts in the 0/−0.7 region and in rhabdomyosarcoma cells in the +2.6/+4.6 Myob-hyperm DMR sequences (Table S4). These two sites might facilitate only weak chromatin interactions when the +2.6/+4.6 sequence is highly methylated and stronger interactions when they are not methylated.

Given that myoblasts have only been used to test genome-wide binding of a very small number of TFs, we also looked for predicted TF binding sites in the cloned regions of *TBX15* from available data for other cell types. Many TFBS were predicted to have binding sites in the Myob-hyperm DMR regions and the promoter region in 5’ end of *TBX15* (Figure 6C, JASPAR database [42]). There were two predicted additional binding sites for MYOD in the +2.6/+4.6 DMR region, for which *in vivo* binding was not seen, as well as multiple sites for STAT3 and MEF2C. In the ChIP-seq Unibind database, binding of various other transcription-stimulatory TFs was seen in the TSS-upstream and TSS-downstream Myob-hyperm DMRs in profiled human cancer cell lines, and endothelial cell cultures, and ESC (Figure 6B; Table S4; Unibind database [39,41]), in which *TBX15* is repressed (Figure 1A). (Table S4). The lack of expression of *TBX15* in ESC can be attributed to the stronger binding of repressive than of transcription-stimulatory proteins at the 5’ end of *TBX15* as seen in the ENCODE database for human ChIP-seq (Figure 6D) and to the related finding of bivalent chromatin in ESC throughout the 5’ end of *TBX15* (Figure 6A and D).

## 3. Discussion

Our study provides evidence that the DNA hypermethylation immediately upstream and downstream of the constitutively unmethylated *TBX15* promoter downmodulates transcription of this gene in primary myoblasts (Figure 7A). These promoter-adjacent DNA sequences were ~10 to 100 times more active in reporter gene assays for promoter or enhancer activity when transfected into myoblasts than when transfected into non-myoblast host cells. This strong activity required demethylation because 86-100% of reporter gene expression was lost upon targeting CpG methylation to the DMR sequences (Figure 4C). Unexpectedly, these DMR sequences exhibited much more reporter gene activity when they were inserted upstream of the vector’s minimal promoter, as is often done (*e.g*., [43]), than when placed downstream of reporter gene. In the downstream position they were only 0.9 kb from the minimal promoter, a favorable distance for enhancer tests [44]. It is likely these promoter-adjacent DMRs (Myob-hyperm DMRs) are part of an extended promoter or context-sensitive enhancer in certain cell/tissue types when unmethylated (Figure 7B). This was demonstrated not only by their demethylation-dependent promoter/enhancer activity but also by *in vivo* correlations between less methylation and more overlap with promoter, enhancer, or open chromatin in *TBX15*-expressing osteoblasts and skin fibroblasts. Our findings argue against the hypothesis that the role of the hypermethylation of these DMRs is to turn off *in cis* an overlapping repressor. A likely explanation for the lack of promoter-border DNA methylation in cells not transcribing *TBX15* is that such cells have no need of fine-tuning *TBX15* expression (Figure 7C).

**Figure 7.**
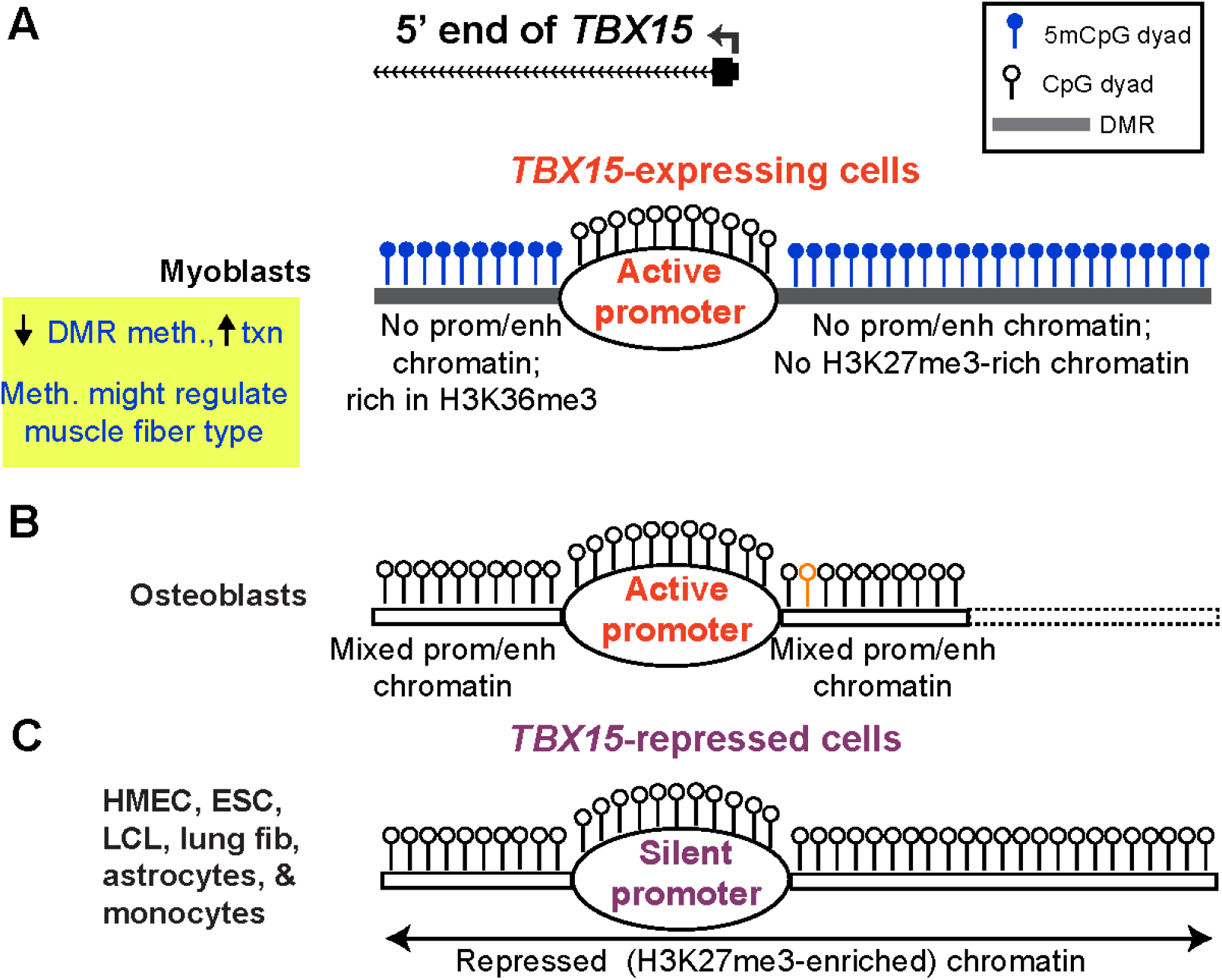
A model for the role of myoblast-hypermethylated DMRs adjacent to the *TBX15* promoter in down-modulating expression of the transcriptionally active gene. (**A**) The high level of methylation of promoter-adjacent DMRs in myoblasts is proposed to prevent their latent enhancer and/or extended promoter activity (enh/prom chromatin) *in vivo* but to allow core promoter activity and downstream enhancer activity (Figures 1 and S8). Certain types of SkM with less methylation at these DMRs may have higher *TBX15* activity due to turning on these promoter-adjacent upregulatory elements; meth, methylation; txn, transcription. (**B**) Some cell types that specifically express *TBX15*, like osteoblasts, have low or no methylation coupled with strong enh/prom chromatin adjacent to the promoter region (Figure S6E). Dotted lines, osteoblast methylome data are from RRBS and so have limited coverage; orange lollipop, a SNP (rs1106529) that is strongly associated with bone mineral density and is located in extended promoter chromatin in osteoblasts overlapping the upstream Myob-hyperm DMR [17]. (**C**) Repressive histone modification (H3K27me3) is seen in many cell and tissue types that have little or no DNA methylation at the DMRs. Such cells with silent *TBX15* alleles do not need DNA hypermethylation-linked fine-tuning of expression.

Consistent with the much higher promoter/enhancer activity of these *TBX15* DMRs in transfected myoblasts than in transfected non-myoblasts, one of these DMRs has a binding site in human myoblasts for MYOD, a central SkM lineage-specific TF, as well as a predicted site for MEF2C (Figure 6), a TF involved in various differentiation processes including myogenesis and repair of SkM [45]. Similarly, in the other *TBX15* DMR used for transfection, there were five predicted MEF2C sites interspersed with three predicted sites for STAT3, a signal transducer and transcription-activating TF involved in muscle satellite cell expansion and SkM repair, among other pathways [46]. We propose that the Myob-hyperm DMRs at the borders of the *TBX15* promoter, when unmethylated, upregulate transcription *in vivo* in certain SkM-lineage cell types. The more distal upstream Myob-hyperm DMRs may be suppressing activity of potential promoters overlapping CpG islands (Figure 1) [47].

We found less than a two-fold increase in *TBX15* RNA in human myotubes relative to their myoblast cell precursors (Table S3) and similar DMRs at the gene in both cell types. Lee *et al*. reported a twelve-fold increase in *Tbx15* RNA when a murine myoblast cell line (C2C12) was induced to differentiate to myotubes [10]. Differences in these results might be due to differences in the protocols used for induction of myoblasts to form myotubes. Differentiation *in vitro* of mononuclear myoblasts to elongated, broadened multinucleated myotubes involves removal of fetal bovine serum from the culture medium. The larger increase in *TBX15* RNA levels upon differentiation in the study of Lee *et al*. than in ours might be due to our less severe differentiation protocol in which fetal bovine serum is replaced with 1.5% horse serum (HS) for only one day followed by 4 d with 15% HS rather than the more standard procedure of Lee *et al*. that uses 2% HS for 4 d [10].

The enhancer and promoter chromatin at the 5’ end of *TBX15* in osteoblasts (Figure 7B) is part of a super-enhancer that extends for ›10 kb downstream of the TSS [14,17]. Super-enhancers strongly upregulate gene expression and are seen most frequently at developmental genes [33]. We previously reported that this super-enhancer contains a SNP (rs1106529; orange lollipop, Figures 7B and S6C) which is strongly associated with bone mineral density (and obesity-risk traits) [17]. This SNP is in the Myob-hyperm DMR immediately upstream of the promoter region, in DNA sequences that are unmethylated in osteoblasts (Figure S6D). It is not in linkage disequilibrium (r^2^ > 0.2) with any other SNP, which would have complicated the determination of its biological importance [17]. Therefore, we propose that osteoblasts are an example of a cell type that benefits from DNA methylation-sensitive upregulation of *TBX15* by enhancer/promoter chromatin bordering the active promoter of *TBX15* and overlapping a Myob-hyperm DMR.

Differences in epigenomic profiles around the *TBX15* promoter in skin fibroblast cell strains from different body sites suggest differential regulation of this gene in skin. The need for fine-tuning expression of *TBX15* in skin is evidenced by dynamic position-dependent differences in *Tbx15* expression in dermal cells during mouse embryogenesis that contribute to mouse coat patterning [48]. In a genome-wide study of human adipocytes, Bradford *et al*. [37] reported several subcutaneous *vs*. visceral adipocyte DMRs that they described as close to the 5’ end of *TBX15* among the 2108 DMRs that they identified. These *TBX15* DMRs exhibited a positive correlation between hypermethylation and preferential expression in subcutaneous adipocytes compared with matched visceral adipocytes. We localized their 5’ *TBX15* adipocyte DMRs to the borders of the *TBX15* promoter region and found that they overlap Myob-hyperm DMRs (Figure S2). Bradford and coworkers suggested that the function of this promoter-bordering DNA hypermethylation is to prevent spreading of repressive chromatin into the active promoter in subcutaneous adipocytes. However, our results favor the hypothesis that this *TBX15* promoter-adjacent DNA hypermethylation in both subcutaneous adipocytes and myoblasts prevents the formation of promoter-adjacent enhancer chromatin that would lead to high levels of transcription of *TBX15* in these cells.

In another study of *TBX15* in adipose cells, Ejarque *et al*. [49] examined the methylomes of human adipose-derived stromal/stem cells from subcutaneous adipose tissue derived from either obese or lean middle-aged females. Cells from lean individuals had approximately 2.5-fold more *TBX15* RNA than the analogous cells from the same body depot in obese individuals. They showed that upregulation of *TBX15* correlated with less methylation in the regions we identified as promoter-adjacent Myob-hyperm DMRs, a finding that is consistent with our model (Figure 7A and B). However, in that study DNA methylation changes were more moderate and seem to be behaving like a continuously adjustable, rather than an on/off, regulator.

TBX15 can act as both a transcription repressor and activator [50,51]. It can downregulate mitochondrial oxidation rates in conjunction with Ampk phosphorylation in SkM and myoblasts, and high *Tbx15* expression in a subfraction of murine subcutaneous adipocytes correlates with lower levels of oxidative metabolism markers [6,10]. In C2C12 myoblasts and murine embryos, Tbx15 was implicated in indirectly upregulating *Igf2*, which controls embryonic myogenesis [10]. Tbx15 induced proliferation of mesenchymal precursor cells and prehypertrophic chondrocytes, but only transiently during embryogenesis, as concluded from studies of a *Tbx15* null mutant *vs*. normal mouse embryos [2]. In cancer cell lines, human *TBX15* was shown to have an anti-apoptotic function that could be partly mediated by its suppression of transcription of several apoptosis-associated *BCL2* family genes [4,52]. Therefore, while there is much more to be learned about TBX15’s cell type-specific regulation of transcription, clearly it can play pivotal roles in differentiation, homeostasis, and changes in cell physiology, which may necessitate careful modulation of its transcription levels partly by promoter-adjacent DNA hypermethylation.

From an examination of human RRBS methylomes, we previously reported that T-box genes are overrepresented among the genes with myoblast DNA hypermethylation [30]. In the current, much more extensive WGBS/EM-seq study, we found that *TBX18, TBX2, TBX3*, and *TBX1*, like *TBX15*, exhibited Myob-hyperm DMRs bordering the active promoter. In contrast *TBX4, TBX5*, and *TBX20* displayed this myoblast DNA hypermethylation bordering on and encroaching into the silent H3K27me3-enriched promoter region (Figures 1 and S3 - S5). The high density of CpGs around or in the promoter regions of these eight T-box genes is unlike that of most tissue-specific genes but is found at a higher frequency in genes encoding tissue-specific TFs [53]. Although we saw evidence of frequent silencing of these T-box genes in cancer cell lines by both polycomb repressed chromatin (H3K27me3) and DNA hypermethylation (Figure S6 and data not shown [23]), one cancer cell line, U87 astrocytoma cells, expressed *TBX15* at moderate levels [38]. In contrast, astrocytes have negligible expression of this gene. Importantly, U87 cells were the only cancer cell line in the RRBS database at the UCSC Genome Browser [23,31] displaying myoblast-like hypermethylated DMRs around an unmethylated promoter region (Figure S6). These findings suggest the acquisition of expression-linked promoter-border DNA hypermethylation in certain cancers, which might contribute to carcinogenesis through antiapoptotic effects of *TBX15* upregulation [4].

Similar to myoblasts, two SkM muscle samples for which there are available WGBS profiles, displayed *TBX15* DMRs similar to the Myob-hyperm DMRs although with less extensive methylation (Figures 1 and 5). These SkM samples were both psoas muscle. However, two other SkM samples of unknown muscle type were largely unmethylated around the *TBX15* promoter (Figure 5). Analyses of epigenomics and transcriptomics in SkM are complicated by many factors including cell heterogeneity, muscle fiber type composition differences and other SkM subtype differences and can be influenced by exercise, muscle disuse, aging, gender, and diet [54–60]. In mice, scRNA-seq indicated that only 68% of the nuclei in soleus and quadriceps are the myocyte nuclei [61]. Varying proportions of non-myocyte cells can be found in SkM tissues from different parts of the body [57]. Despite these complicating factors, murine *Tbx15* was found to be predominantly expressed in SkM types enriched in fast-twitch glycolytic muscle fibers rather than in slow-twitch myofibers or fast oxidative myofibers [10], and its expression has been used as a marker of fast glycolytic muscle fibers [62]. Different muscle types are mixtures of slow and fast myofibers, including some hybrid slow/fast myofibers. Lee *et al*. [10] reported and Terry *et al*. [57] confirmed that mice had about twice as high *Tbx15* RNA levels in gastrocnemius, tibialis anterior, and extensor digitorum longus muscle (all of which are enriched in glycolytic myofibers) relative to soleus muscle, which has a higher percentage of oxidative myofibers.

Human psoas SkM is mostly a body-support muscle. Accordingly, it has a low content of fast glycolytic myofibers [63]. Psoas SkM exhibited less enhancer/extended-promoter chromatin at the 5’ end of *TBX15* than did two leg muscle samples for which chromatin epigenomic profiles, but not methylomes, were available (Figure 5D). Although the donors for the leg muscle samples (54 y M and 72 y F) were much older than those for the examined psoas sample (mixed 3-y M and 34-y M), we favor the explanation that the observed chromatin epigenetic differences between the psoas and leg samples at *TBX15’s* 5’ end resulted from differences in myofiber composition rather than age differences. A meta-analysis of 908 SkM samples did not identify *TBX15* among the genes with age-associated DMRs [58]. Furthermore, fast-twitch glycolytic fibers in humans, which in mice are associated with high *Tbx15* expression [10], have been found to decrease, rather than increase, with age [55]. We propose that the absence of *TBX15* promoter-adjacent DNA hypermethylation in two RRBS-analyzed SkM samples from unspecified parts of the body (Figure 5C) is due to their derivation from muscle types with especially high expression of *TBX15*. Because myofiber type plays major and complex roles in muscle performance, muscle formation, muscle repair, and sarcopenia [64,65], the epigenetic regulation of postnatal *TBX15* expression is likely to be important for normal muscle function and maintenance.

*TBX15* exhibits higher expression in both postnatal SkM (gastrocnemius, a lower leg muscle) and fetal human SkM than in other examined tissues and is expressed in both SkM myocytes and muscle satellite cells (Tables S1 and S2). Nonetheless, the major phenotype of homozygous loss of function of *TBX15* in humans (Cousin syndrome) or in mouse knockout models is major deformities in the skeletal system reflecting a critical role for its encoded protein in embryonic skeletal bone formation [2,66]. In a mouse strain with homozygous loss-of-function of *Tbx15*, a few changes in the musculature were noted [3,67]. The major limb malformations in humans or mice associated with loss of TBX15 activity could largely mask muscular defects given the interrelations of SkM and bone functionality. Interestingly, a T-box gene called *Tbx15/18/22* in *Ciona intestinalis*, an aquatic invertebrate, is essential for normal transcription of many muscle structural genes [68]. Of all the human T-box genes, only *TBX15* has strong specificity for fetal human SkM cells (Table S2).

The biological importance of epigenetic fine-tuning of *TBX15* transcription may be related to the phenotypic effects of partial loss of T-box TFs, in general. Deleterious mutations in all but one (*TBX18*) of the 17 T-box family genes cause disease phenotypes or prenatal lethality when homozygous in humans, and mutations in 10 of the genes also give a phenotype when heterozygous [3]. Such heterozygosity usually decreases the wild-type levels of the corresponding gene product approximately two-fold. Therefore, this type of heterozygote phenotype is evidence for the importance of close regulation of expression of T-box genes. Although *TBX15* heterozygosity for loss-of-function mutations in humans and mice has not been reported to confer overt skeletal abnormalities [3,9], careful examination of heterozygous knock-out mice revealed SkM [10] and facial phenotypic differences from wild-type mice [69]. Lee *et al*. [10] found that these heterozygotes, which had a ~40% decrease in *TBX15* mRNA and protein, had a significant (~10%) decrease in muscle mass in tibialis anterior, a type of muscle consisting predominantly of glycolytic fibers, that was not seen in soleus muscle, a mostly oxidative type of muscle. The analogous homozygotes had a yet larger decrease in muscle mass (~25%) due to changes in muscles enriched in glycolytic myofibers. Our results suggest that large increases in DNA methylation can repress or maintain repression of enhancer-like activity surrounding the unmethylated *TBX15* promoter in muscle fibers. Furthermore, graded changes in this methylation might give partial enhancer-like activity to these promoter-adjacent regions. In both cases, promoter border hypermethylation in *TBX15* is ideally suited for the dynamic regulation of muscle physiology.

## 4. Materials and Methods

### 4.1 Preparation of DNA constructs, transfection, and in vitro DNA methylation

Reporter gene constructs were prepared by overlap extension PCR (Table S5) or by using the Gibson assembly kit (NEBuilder HiFi Assembly, New England Biolabs) as previously described [28]. The vectors (InvivoGen/Invitrogen) were pCpGfree-Lucia or pCpGfree-promoter-Lucia (Vectors 1 and 2, with or without a human EF-1α-derived minimal promoter, respectively, Figure 3). These vectors have a Lucia luciferase reporter gene and no CpGs. The inserts for cloning were obtained by PCR on mixed human brain and placenta DNAs using the primers shown in Table S5. Recombinant plasmid structure was checked by partial DNA sequencing and restriction site analysis. Transfection into C2C12 or MCF-7 cells utilized a lipid-based reagent (Fast-forward protocol, Effectene reagent, Qiagen). As a reference for transfection efficiency, pCMV-CLuc 2 (New England Biolabs) encoding the Cypridina luciferase was co-transfected with the test construct. About 48 h after the transfection, Lucia and Cypridina luciferase activity was quantified by bioluminescence from aliquots of the cell supernatant (BioLux Cypridina Luciferase assay kit, New England Biolabs; Quanti-Luc, InvivoGen). Reference plasmid-normalized luciferase activity was from the average of three independent transfections. Methylation of the plasmids was targeted just to the *TBX15* inserts, which were the only CpG-containing sequences, by incubating the DNA construct (1 μg) with 4 units of SssI methylase and 160 μM S-adenosylmethionine (New England Biolabs) for 4 h at 37°C or mock-methylating by incubating in the absence of S-adenosylmethionine. A similar plasmid construct that contained three BstUI CGCG sites was methylated as above and shown thereafter to be fully resistant to BstUI cleavage.

### 4.2 EM-seq and WGBS on myoblast DNA and determination of DMRs and LMRs

The myoblasts used for DNA isolation were non-transformed cultures derived from quadriceps biopsies of control individuals [70]. Although primary myoblasts, especially from commercial sources, are often contaminated with large numbers of fibroblast-like cells, which can provide misleading results in DNA methylation analyses, we demonstrated that all of our batches of myoblasts contained >90% desmin-positive cells. WGBS of myoblast cell line Myoblast 3 [30] was performed by standard methods [71]. Methylation profiling by EM-seq of Myoblast 3 and two additional myoblast cell strains (Myoblasts 1 and 6) was done as previously described [72]. This involved the enzymatic oxidation of 5mC (TET2) to 5hmC residues and then to 5-carboxylcytosine (5caC) residues followed by glucosylation (T4-phage β-glucosyltransferase) of any remaining 5hmC, conversion of C residues to U residues (APOBEC3A), and PCR [24]. In brief, 0.2 μg of DNA was used for EM-seq library preparation using the NEBNext^®^ Enzymatic Methyl-seq kit for Myoblasts 1, 3 and 6 in duplicate. The resultant libraries were cleaned (NEBNext^®^ sample selection beads) and duplicates were pooled. The final library pool was diluted to 1.5 nM for NovaSeq (illumina) sequencing.

For determining myoblast DMRs, the EM-seq data for the three examined myoblast cell strains were compared to WGBS profiles of foreskin fibroblasts (Skin Fib 2) [73], adipose-derived mesenchymal stem cells induced to differentiate to adipocytes [36], prostate epithelial cells [74], human mammary epithelial cells (HMEC [75]), prenatal lung fibroblasts (IMR90) and ESC [76]; the last three were cell lines established from non-malignant cells and the others are cell strains. We had previously determined SkM (psoas) DMRs by comparing WGBS profiles [19] from psoas to those of heart (left ventricle), aorta, monocytes, lung, and subcutaneous adipose tissue [22,77,78]. To verify that differences were not associated with technical effects, EM-seq myoblast methylation profiles were initially compared to a WGBS methylation profile from one of the three cell strains. While there was significant biological variation among the three cell strains, results indicated that the proportion of differentially methylated sites between the EM-seq and WGBS profiles for the same cell line was consistent with random variation (data not shown). DMRs between the three EM-seq profiles and the group of five cell cultures were determined using a two-phase process, with significantly differentially methylated sites identified via generalized linear models and aggregated into DMRs based on the Uniform Product distribution for p-values as previously described [78]. Low methylated regions (LMRs) shown in the figures refer to regions with significantly lower DNA methylation than in the rest of the same genome as determined using the method of Song *et al*. [21].

### 4.3 Bioinformatics

Most of the bioinformatic profiles were from the UCSC Genome Browser using the hg19 (mainly) or hg38 reference genomes and are shown with hg19 coordinates in the figures [23]. RefSeq Curated gene isoforms are shown unless otherwise specified. WGBS profiles of genome-wide CpG methylation of tissues and cell cultures other than myoblasts (see above) were used for most DNA methylation comparisons and were complemented with RRBS data for *TBX15* methylation, including some previously described cell or tissue profiles [14,31]. Unless otherwise stated, the SkM sample for WGBS was a mixture of psoas DNA from a 3-y male and a 34-y male. Human transcription data was from the following UCSC Browser tracks or hubs: cultured cells (strand-specific RNA-seq, ENCODE/Cold Spring Harbor Lab, or non-strand specific RNA-seq, Transcription Levels Assayed by RNA-seq on 9 cell lines/ENCODE [15]); GTEx (medium TPM from RNA-seq from hundreds of samples for each tissue [35]; Table S1), nine tissues (Burge Lab RNA-seq [79]), Fetal Gene Atlas binned by organ or cell type from Cao *et al*. (scRNA-seq on samples from ~72 to 129 days post-conception fetuses [80]; Table S2), Muscle RNA binned by biosample from De Micheli *et al*. (scRNA-seq on postnatal muscle tissues [54]; Table S2); RNA-seq analysis of multiple types of SkM shown in the hg38 reference genome [19,23], and 5’ Cap Analysis of Gene Expression (CAGE; RIKEN Omics Science Center [81]. The quantitation of cell culture-derived RNA-seq data for poly(A)^+^ RNA was previously described [30]. In addition, for extensive scRNA-seq, the poly(A)^+^ RNA Human Protein Atlas (scRNA-seq on tissues [38]; Table S1; Figure S1) was employed. For comparisons of RNA levels in myoblasts and myotubes, we used previously generated RNA-seq data for poly(A)^+^ RNA from myoblast cell strains from our lab [82]. CGI identification followed the definitions at the UCSC Genome Browser with islands between only 200 and 300 bp identified by their light green color in figures.

The 18-state chromatin state segmentation analysis (chromHMM, AuxilliaryHMM, Roadmap Epigenomics [19]) was used for determination of chromatin states. The color coding in figures of gene regions is as follows: red, promoter or mixed promoter/enhancer chromatin (States 1 - 4); light or dark green, H3K36me3-enriched chromatin (States 5 and 6); orange or yellow-green, enhancer chromatin (States 7 - 10); light yellow, weak enhancer chromatin (State 11); H3K9me3- and H3K36me3-enriched chromatin and ZNF-gene associated chromatin (State 12); blue, H3K9me3-associated (State 13); reddish brown or gray-green, bivalent poised promoter and enhancer (States 14 and 15, respectively); light or dark gray, H3K27me3-associated repressed chromatin (States 16 and 17); and white, low signal for H3K27ac, H3K27me3, acetylation or methylation, H3K4me1, H3K4me3, or H3K9me3 (State 18). Also available at the UCSC Genome Browser [23] are DNase-seq profiles of various skin fibroblast cell strains (Roadmap Epigenetics Project for Figure 2 and ENCODE project/University of Washington for Figure 5), CTCF binding profiles for cell cultures (ENCODE), predicted TFBS from the JASPAR database [42], ChIP-seq profiles combined with TFBS prediction from UniBind [39], and ChIP-seq profiles from ENCODE 3 Transcription Factor ChIP-seq Clusters. The presence of super-enhancers was determined using the SEdb tool [34] although the sizes of the super-enhancers were determined by visual examination of H3K27ac and H3K4me1 tracks (vertical viewing range, 0 – 10).

## 5. Conclusions

DNA methylation near the promoter region is usually associated with silenced genes. In contrast, *TBX15* had strong hypermethylation bordering its unmethylated CGI-containing promoter in myoblasts and psoas skeletal muscle that correlates with its preferential expression in skeletal muscle. Results from our reporter gene assays and bioinformatic comparisons of many cell and tissue types indicate that DNA hypermethylation at the upstream and downstream borders of the *TBX15* promoter helps to prevent overexpression of *TBX15*. Previous functions suggested for this *TBX15* promoter-adjacent hypermethylation were silencing of putative repressor elements, protecting against the expansion of nearby repressed chromatin into the promoter, or directing the use of alternate promoters. Our results suggest that the loss of this methylation or a decrease in the extent of methylation *in vivo* is associated with higher expression of *TBX15* by removing repression from the silenced enhancer-like DNA sequence elements in these hypermethylated DMRs. Our results also present a cautionary tale about how cancer-related DNA hypermethylation at a CpG island near the 5’ end of a gene need not correlate with silencing of the gene. While high DNA methylation levels within a CpG-rich promoter are well known to strongly repress transcription, such methylation in promoter-adjacent regions, may only downmodulate, rather than silence expression and may not be found in cell types in which the gene is otherwise silenced by repressive chromatin. These findings reinforce the importance of understanding the genetic and chromatin context of regions being examined for DNA hypermethylation associated with differentiation, physiological changes, or disease.

## Supporting information

Promoter-adjacent DNA hypermethylation can downmodulate gene expression: TBX15 in the muscle lineage-Supplemental figures

Promoter-adjacent DNA hypermethylation can downmodulate gene expression: TBX15 in the muscle lineage-supplemental Tables

## Supplementary Materials

Figure S1: *TBX15* is preferentially expressed in myocytes, fibroblasts, and smooth muscle cells in skeletal muscle (SkM) tissue. Figure S2: Whole-genome methylome profiles of *TBX15* for the three myoblast cell strains and six non-myogenic cell cultures used for determining myoblast DMRs: Similarities of myoblast *vs*. non-myoblast DMRS (this study) to subcutaneous *vs*. visceral adipocyte DMRs from Bradford *et al*. 2019. Figure S3: Myoblast-hypermethylated DMRs adjacent to promoters in *TBX1* and *TBX2*, which are preferentially expressed in myoblasts. Figure S4: Myoblast-hypermethylated DMRs surround the promoter in *TBX3*, which is preferentially expressed in myoblasts but overlap the promoter of *TBX20*, which is repressed in myoblasts. Figure S5: Myoblast-hypermethylated DMRs overlap promoters of *TBX4* and *TBX5*, which are repressed in myoblasts. Figure S6: The intergenic region between *TBX15* (cell-type specific expression) and *WARS2* (broad expression) displays multiple myoblast-hypermethylated DMRs but less osteoblast hypermethylation. Figure S7: *TBX15* and *TBX1* but not *TBX15’s* neighbor *WARS2* display more expression in female skeletal muscle than in male skeletal muscle. Figure S8: *TBX15* is the only gene preferentially expressed in the skeletal muscle lineage in its gene neighborhood but there is a very far downstream enhancer chromatin region overlapping a hypomethylated DMR in skeletal muscle. Table S1: RNA-seq profiles showing that *TBX15* is the T-box gene with the strongest preference for expression in both skeletal muscle myocytes and skeletal muscle tissues. Table S2: Expression of *TBX15* among human fetal organs and cells and postnatal skeletal muscle from different anatomical sites. Table S3: Cell type-specificity of RNA levels from T-box family genes among different types of human primary cell cultures. Table S4: Transcription factor binding sites demonstrated for various cell types in the 5’ end of *TBX15*. Table S5. Oligonucleotides used for fusion cloning of 5’ *TBX15* sequences into CpG-free Luciferase reporter vectors.

## Author Contributions

KCE and ME conceived the study, made the reporter constructs, did the transfection experiments and wrote the manuscript. ML determined the DMRs for myoblasts and CB determined the LMRs from myoblast methylomes generated by SS and POE, who were under the direction of SP. The myoblast DNA was previously isolated from primary cells grown and characterized by immunocytochemistry under the direction of ME.

## Data Availability Statement

Data is contained within the article and supplementary material.

