## Supplementary material for "Promoter-adjacent DNA hypermethylation can downmodulate gene expression: *TBX15* in the muscle lineage": Promoter-adjacent DNA hypermethylation can downmodulate gene expression: TBX15 in the muscle lineage-Supplemental figures

**Supplementary Figures S1 – S8 for**

**Promoter-adjacent DNA hypermethylation can downmodulate gene expression:**

***TBX15* in the muscle lineage**

**Kenneth C. Ehrlich, Michelle Lacey, Carl Baribault, Sagnik Sen, Pierre Olivier Esteve,**

**Sriharsa Pradhan, and Melanie Ehrlich**

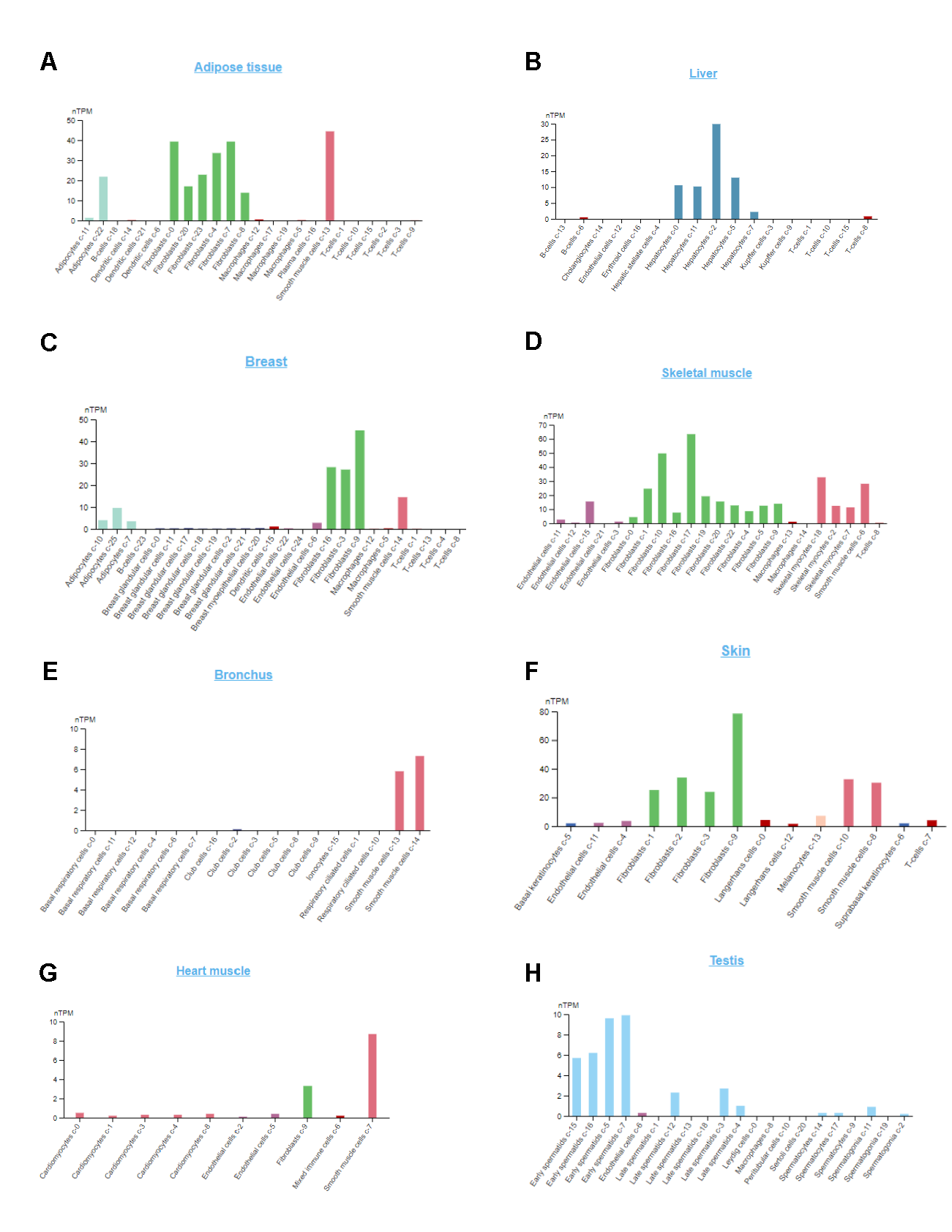

**Figure S1. *TBX15* is preferentially expressed in myocytes, fibroblasts, and smooth muscle cells in skeletal muscle (SkM) tissue.** **A** – **H**. scRNA-seq profiles of *TBX15* RNA in the indicated tissues (Human Protein Atlas [1]) show preferential expression of this gene in myocytes in SkM and in fibroblasts in several tissues, including SkM. There is also preferential expression in smooth muscle cells, early spermatids, adipose tissue, and breast tissue. The transcripts in liver hepatocytes are probably derived from a far downstream promoter and exon 1 [2].

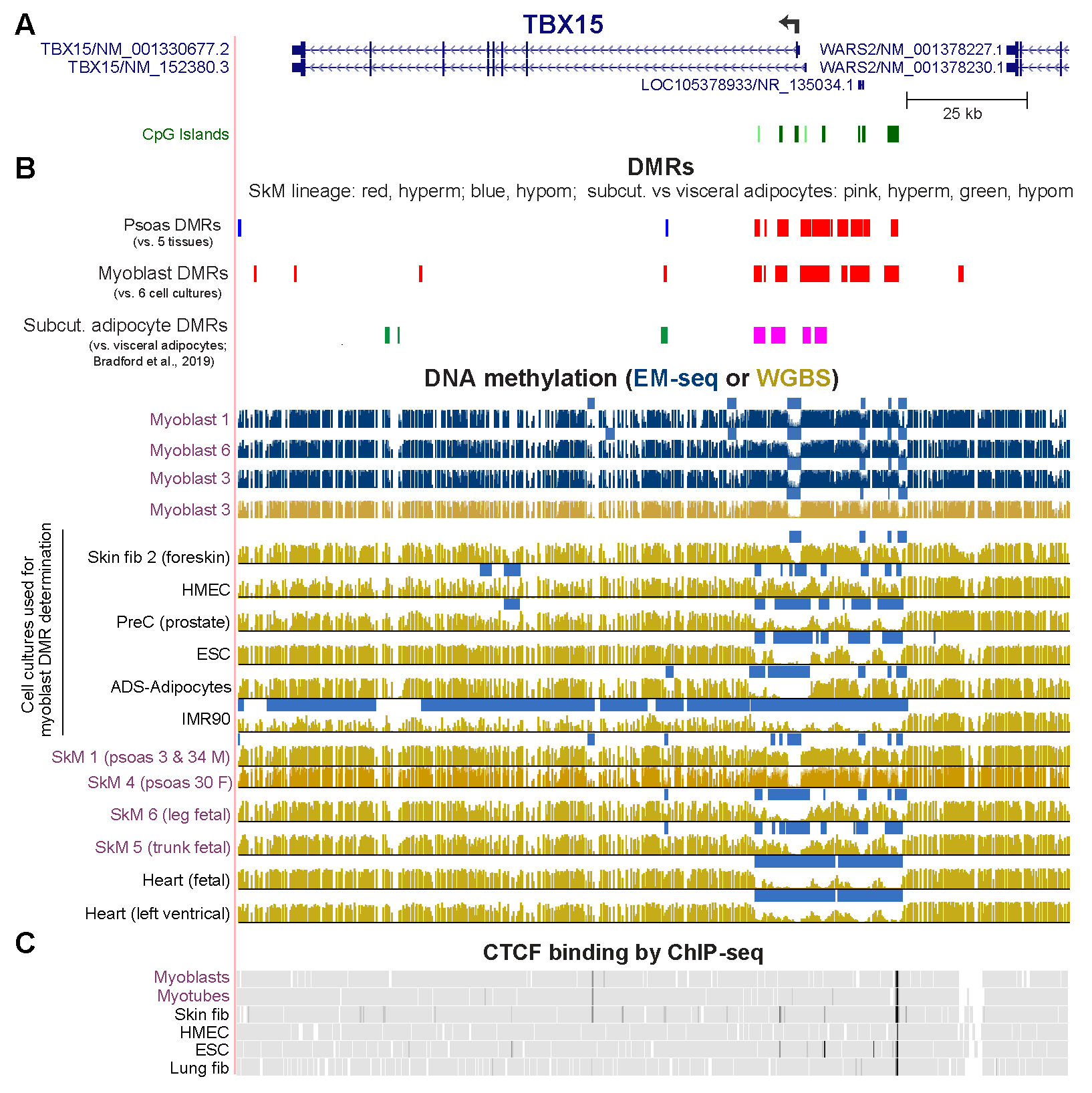

**Figure S2.** **Whole-genome methylome profiles of *TBX15* for the three myoblast cell strains and six non-myogenic cell cultures used for determining myoblast DMRs: Similarities of myoblast *vs.* non-myoblast DMRs (this study) to subcutaneous *vs.* visceral adipocyte DMRs from Bradford *et al*. 2019. A.** *TBX15* and the 3’ end of broadly expressed *WARS2* (chr1:119,414,444-119,586,756, hg19; all tracks are aligned). **B.** DNA methylation analyses. Our previously determined SkM (psoas muscle *vs*. five non-muscle tissues) DMRs [3-5], myoblast *vs.* 6 non-muscle cell culture DMRs from this study, and subcutaneous (subcut.) adipocyte *vs.* visceral adipocyte DMRs from a report by Bradford *et al.* [6] with the indicated color coding for hypermethylated (hyperm) and hypomethylated (hypom) DMRs. DNA methylation profiles (as in Figures 1 and 2) with light blue bars indicating low methylated regions and the extent of methylation at each CpG site (full-length bars show 100% methylation). **C**. CTCF binding profiles; only one constitutive CTCF site (present in all examined cell types) is seen. HMEC, mammary epithelial cells; PreC, primary prostate epithelial cell culture; ESC, H1 embryonic stem cells; IMR90, fetal lung fibroblast cell line; fib, fibroblast cell strain; ADS-Adipocytes, subcutaneous adipose-derived mesenchymal stem/stromal cells induced to form adipocytes. Note that ADS-Adipocytes are not the same as *in vivo*-derived adipocytes in the study by Bradford *et al.* [6].

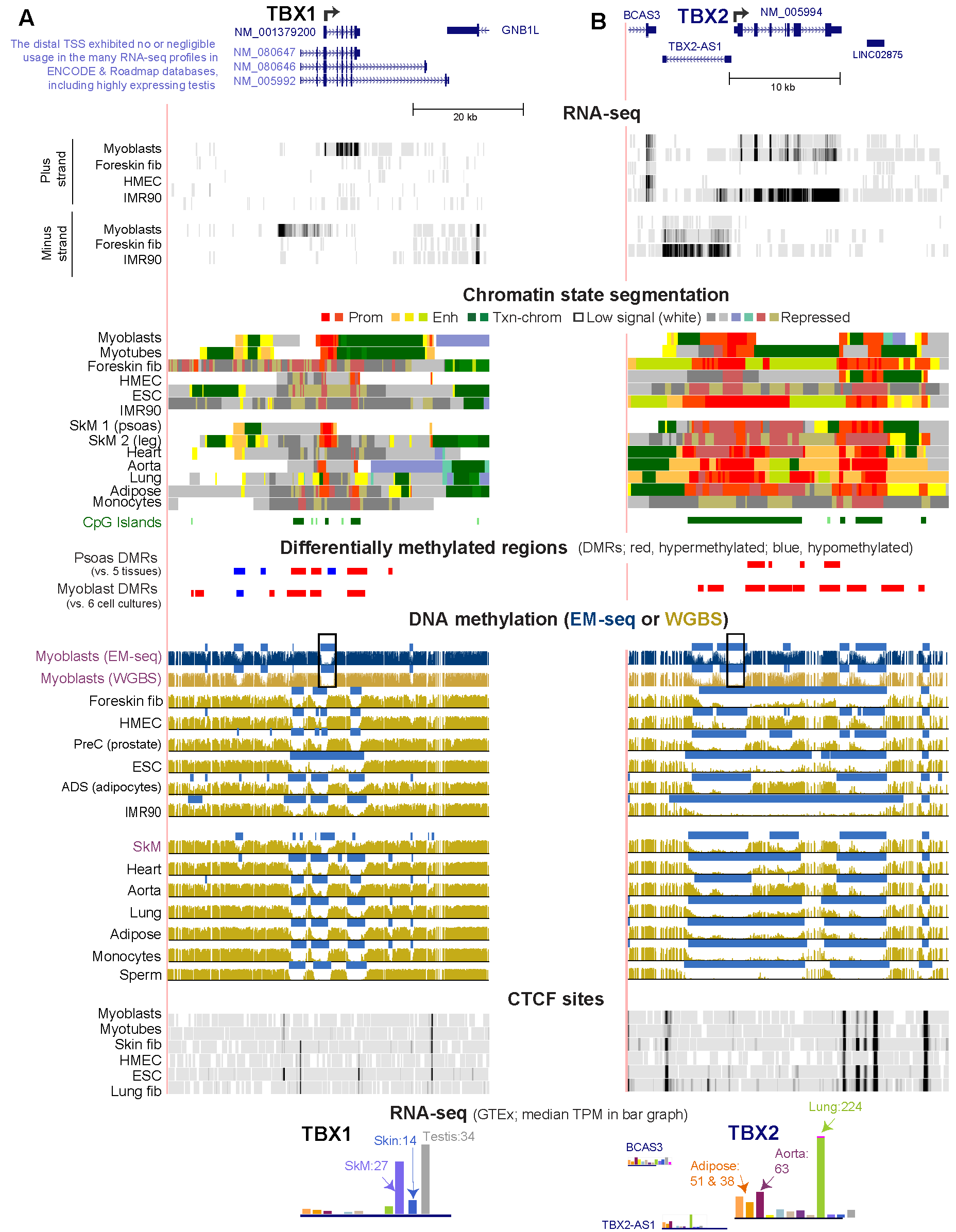

**Figure S3. Myoblast-hypermethylated DMRs surround the main promoter in *TBX1* and *TBX2*, genes that are preferentially expressed in myoblasts.** **A.** *TBX1* RefSeq isoforms (chr22:19,720,265-19,778,385). **B.** *TBX2* RefSeq structure (chr17:59,467,686-59,496,398). In both panels, strand-specific RNA-seq profiles from myoblasts, skin fibroblasts (fib; Foreskin fib 3), mammary epithelial cells (HMEC), and fetal lung fibroblasts (IMR90) are shown. Color-coded chromatin states for the indicated cell cultures or tissues depict promoter or mixed promoter/enhancer (prom), strong enhancer (enh), or repressed (rep) types of chromatin, transcribed chromatin (txn chrom), and chromatin with a low signal for H3K27ac, H3K27me3, and H3K4 or K9 methylation. Postnatal lung fibroblasts (not shown) and IMR90 (fetal lung fibroblasts) had similar chromatin segmentation profiles. As in Figure S2, this figure shows myoblast and SkM DMRs, WGBS and EM-seq methylomes (regions having significantly lower methylation relative to the same genome indicated by light blue horizontal bars); Myoblast 3 was used for EM-seq and WGBS and Skin fib 2 for WGBS. CTCF binding sites from ChIP-seq are depicted. GTEx RNA-seq expression profiles are given as linear-scale TPM bar graphs with some of the median TPM values for a given tissue indicated; adipose tissue bars show subcutaneous adipose on the left and visceral adipose on the right.

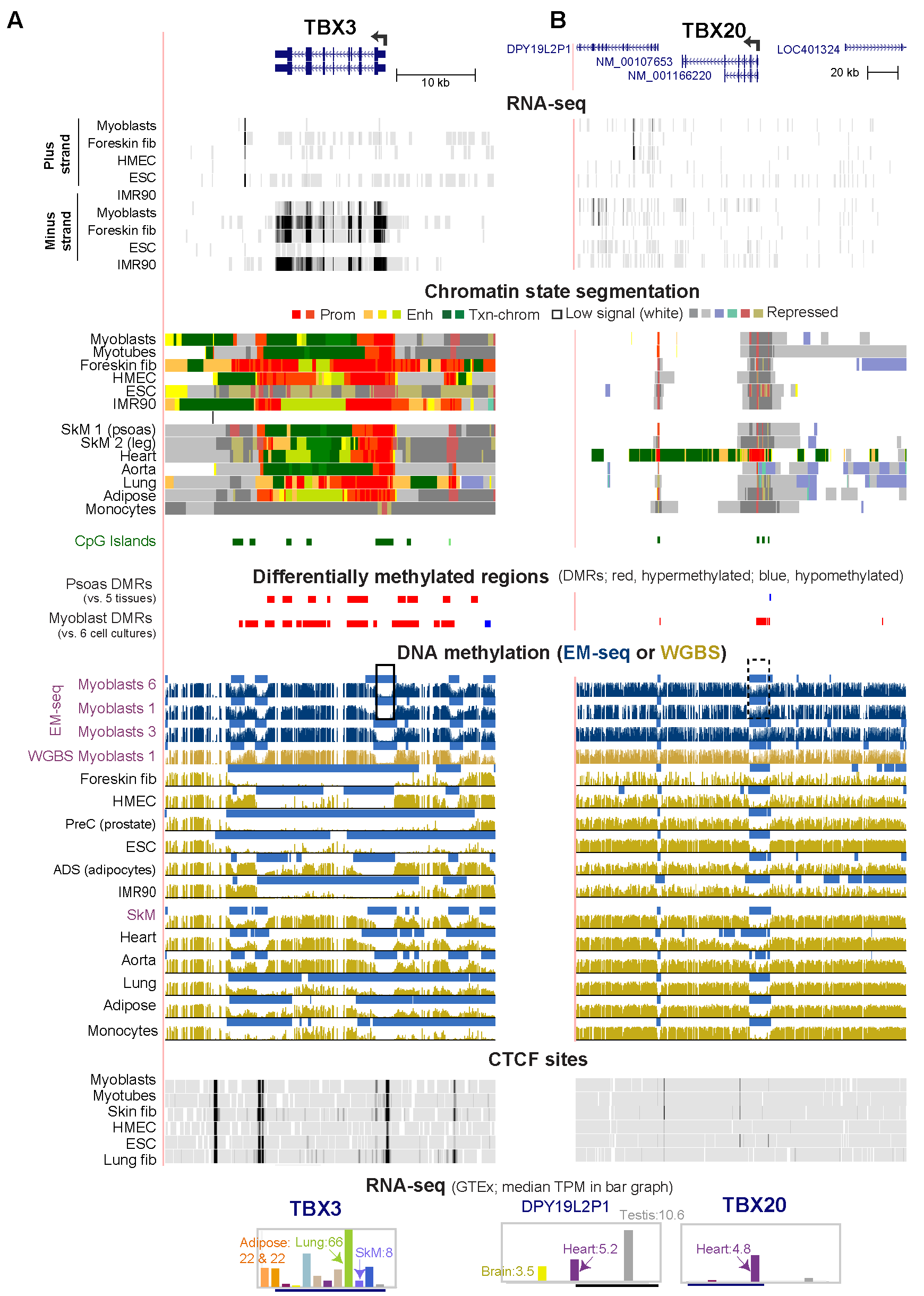

**Figure S4.** **Myoblast-hypermethylated DMRs surround the promoter in *TBX3*, which is preferentially expressed in myoblasts, but overlap the promoter of *TBX20,* which is repressed in myoblasts.** **A.** *TBX3* (chr12:115,094,148-115,135,880) and **B.** *TBX20* RefSeq isoforms (chr7:35,169,412-35,395,411) and transcriptomic and epigenomic profiles are shown as in Figure S3. Postnatal lung fibroblasts (not shown) and IMR90 (fetal lung fibroblasts) had similar chromatin segmentation profiles.

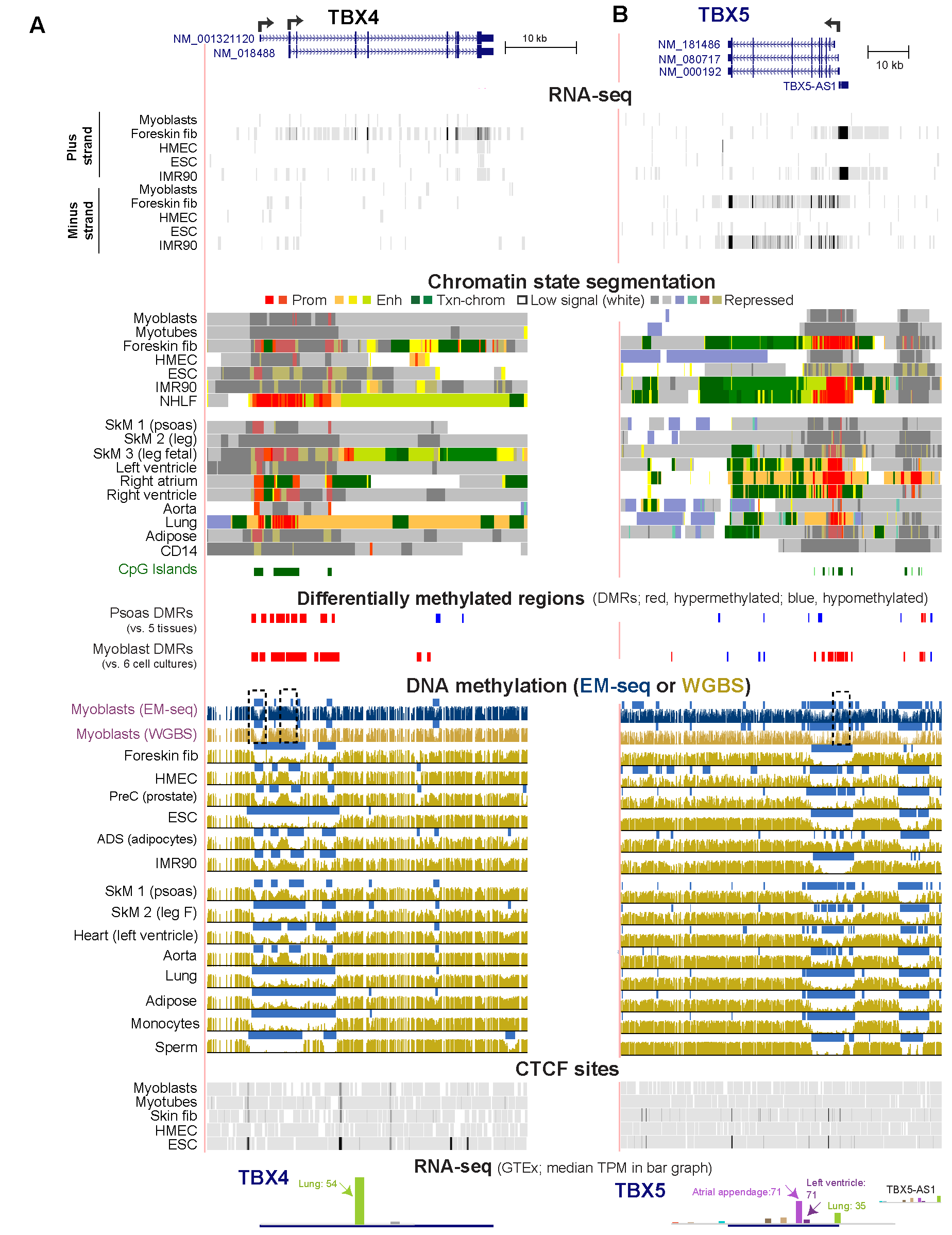

**Figure S5.** **Myoblast-hypermethylated DMRs overlap promoters of *TBX4* and *TBX5,* which are repressed in myoblasts.** **A.** *TBX4* (chr17:59,522,412-59,567,277) and **B.** *TBX5* RefSeq isoforms (chr12:114,739,501-114,896,202) and transcriptomic and epigenomic profiles are shown as in Figure S3.

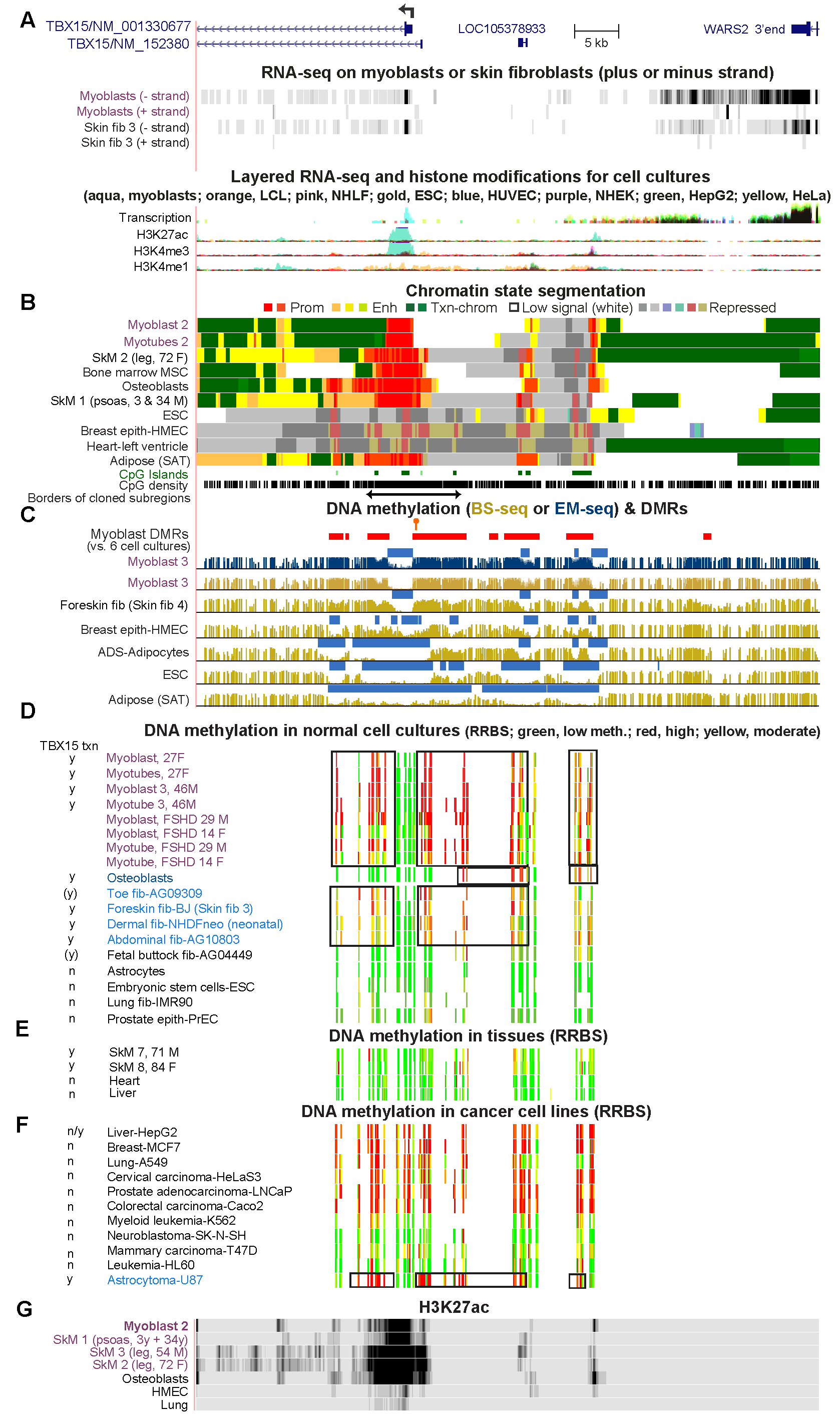

**Figure S6. The intergenic region between *TBX15* (cell-type specific expression) and *WARS2* (broad expression) displays multiple myoblast-hypermethylated DMRs but less methylation in osteoblasts. A**. 5’ end of *TBX15* to 3’ end of broadly expressed *WARS2* (chr1:119,506,614-119,577,071) with strand-specific RNA-seq profiles and layered, color-coded profiles for non-strand specific RNA-seq and H3K27ac, H3K3me3, and H3K4me1 signal profiles in myoblasts (aqua, the only appreciable signal seen at the 5’ end of *TBX15*), a B-cell derived lymphoblastoid cell line (LCL, GM12878, orange), normal human lung fibroblasts (NHLF, pink), H1 embryonic stem cells (ESC, gold), human umbilical vein endothelial cells (HUVEC, blue), epidermal keratinocytes (NHEK, purple), a hepatocarinoma cell line (HepG2, green), cervical cancer cell line (HeLa, yellow); data from the ENCODE Project [7]. The vertical viewing setting for the strand-specific RNA-seq poly(A)^+^ RNA was 0-10 for sensitive detection of poly(A^)+^ RNA signal. **B** and **C.** Chromatin state, CpG islands, and CpG density, the extent of the cloned regions, myoblast DMRs, and methylome tracks, as in previous figures. Orange arrow above a DMR in Panel **C** denotes the SNP (rs1106529) that we previously reported to be strongly associated with bone mineral density [2]. **D.** RRBS DNA methylation data for normal or diseased (FSHD, facioscapulohumeral muscular dystrophy) myoblasts and various other cell cultures not derived from cancers. **E.** RRBS data for normal tissues. **F.** RRBS data for cancer cell lines. **G.** H3K27ac signal profiles (vertical viewing range, 0-10) from the Roadmap Project [8]. Txn, transcription of *TBX15* in the given cell culture or tissue is indicated by y, yes; (y), yes by extrapolation from many similar samples; n, no; or n/y, not from the proximal promoter but from a far downstream liver-specific promoter mentioned in the text; transcription data for panel **G** were from ENCODE profiles at the UCSC Genome Browser or the Human Protein Atlas scRNA database [1].

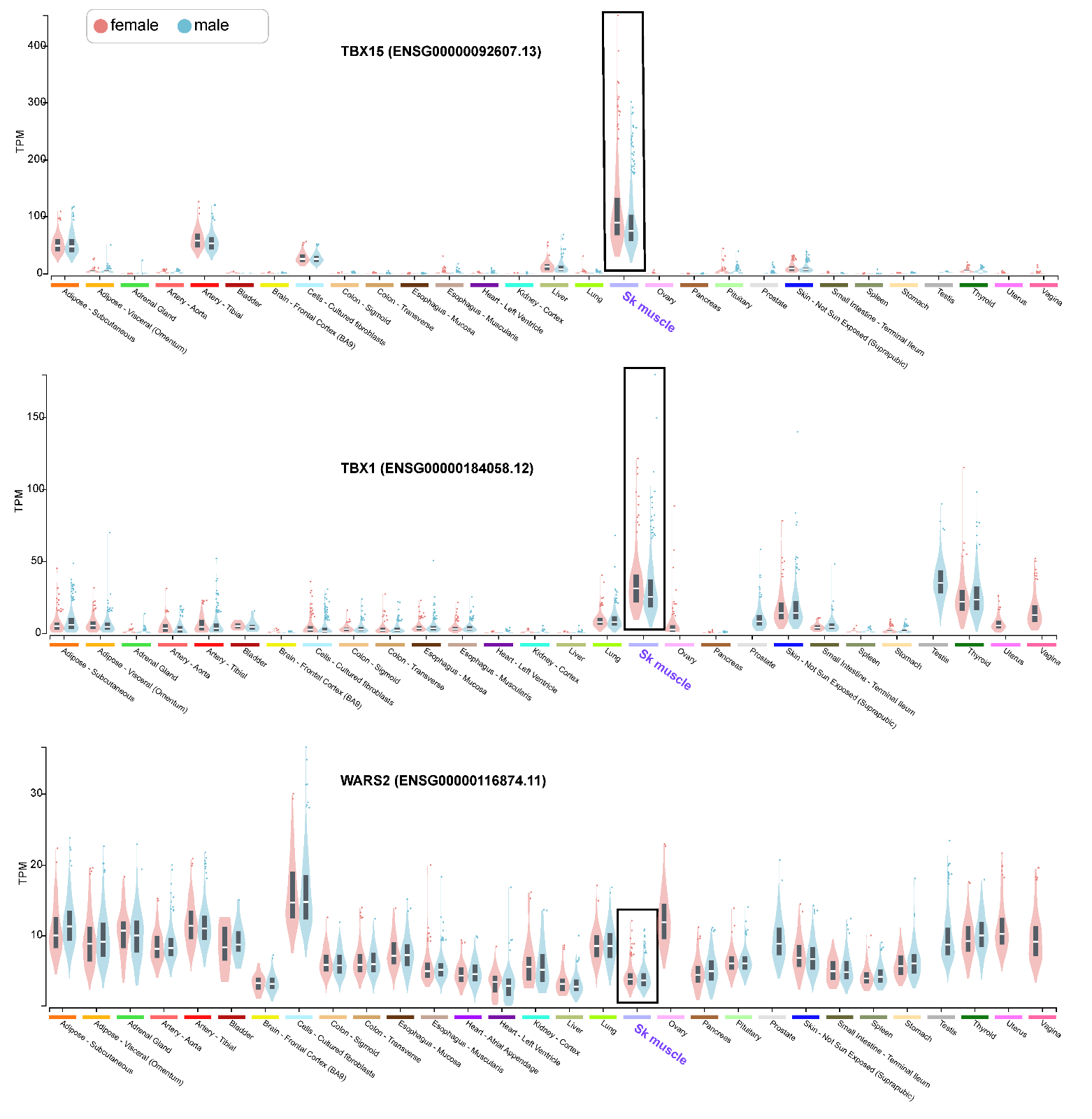

**Figure S7. *TBX15* and *TBX1* but not *WARS2*, (*TBX15*’s neighbor) display more expression in female skeletal muscle than in male skeletal muscle. A – C.** Violin plots of the TPM from hundreds of samples for each tissue type sorted for female *vs.* male expression. The date are from the GTEx database RNA-seq analyses (https://gtexportal.org/), which involved 543 males and 260 females for the SkM samples.

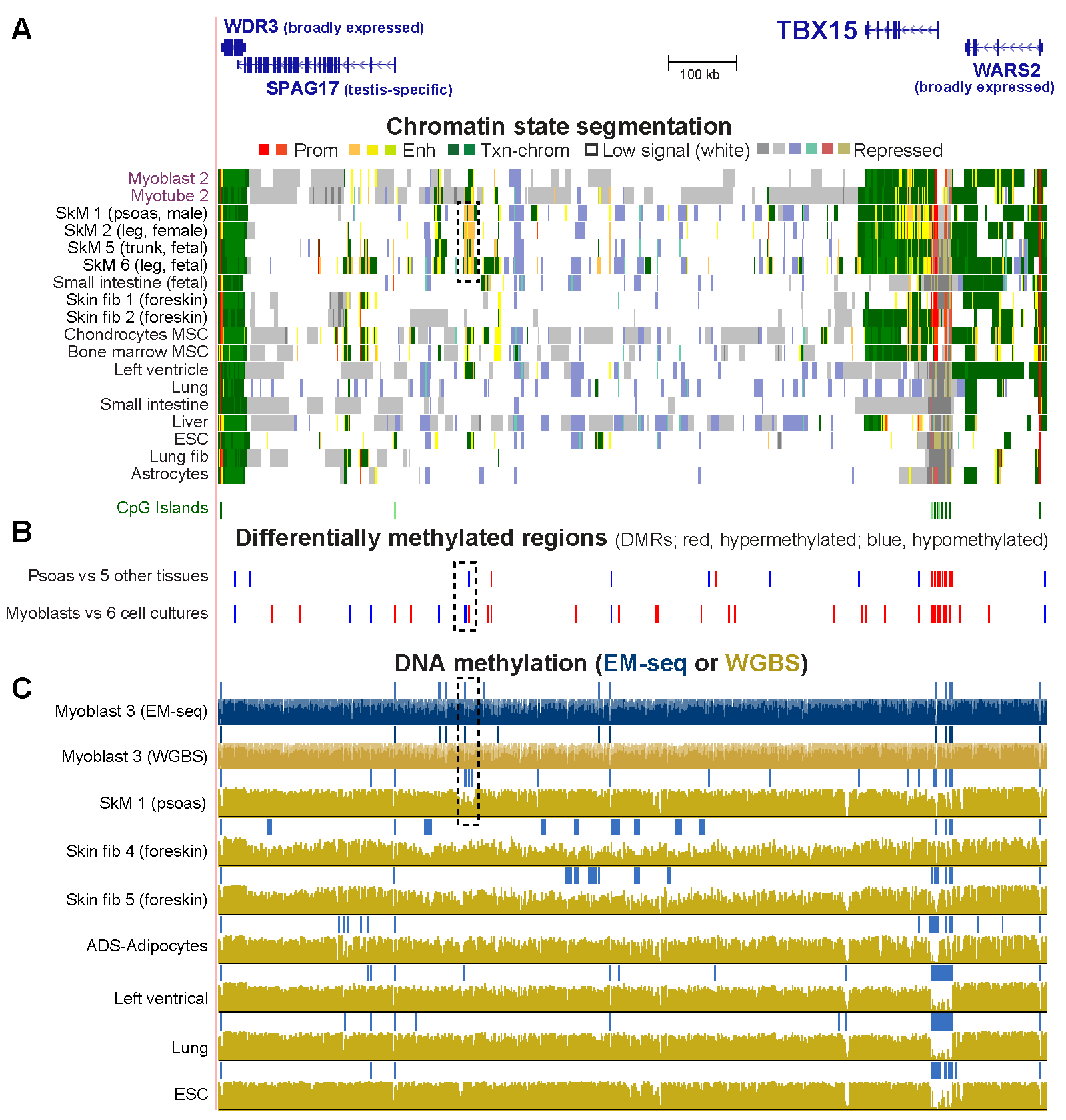

**Figure S8. *TBX15* is the only gene preferentially expressed in the skeletal muscle lineage in its gene neighborhood but there is a very far downstream enhancer chromatin region overlapping a hypomethylated DMR in skeletal muscle. A.** The 1.2-Mb gene neighborhood of *TBX15* (chr1:118,467,598-119,693,423) and chromatin state profiles as in previous figures. **B.** Psoas or myoblast DMRs. **C.** Methylome profiles. The dotted rectangles show enhancer chromatin seen specifically in SkM samples or DNA hypomethylation in myoblasts or, to a larger extent, in SkM.
